# Fatty Acid Oxidation Suppression Reprograms Fibroblasts in Fibrostenotic Crohn’s Disease

**DOI:** 10.64898/2026.05.06.723289

**Authors:** Jihad Aljabban, Ayesh Awad, Benjamin D. McMichael, Valerie Gartner, Viguna Thomas, Benjamin Huan, David Weaver, Grace Lian, Caroline Beasley, Gwen Wan-Jun Lau, Sophie Silverstein, Muneera Kapadia, Anna C. Salvador, Florian Rieder, Jessica E. Thaxton, Terrence S. Furey, Aadra P. Bhatt, Shehzad Z. Sheikh

## Abstract

Fibrostenotic complications represent a major cause of morbidity in Crohn’s disease (CD), yet the cellular mechanisms that drive intestinal fibrosis independent of active inflammation remain poorly understood. Here, we identify impaired fatty acid oxidation (FAO) as a defining metabolic feature of fibroblasts in fibrostenotic CD. Untargeted lipidomics of non-inflamed colonic tissue from CD patients demonstrated enrichment of triacylglycerols and long-chain acylcarnitines, suggesting altered lipid utilization. Across three independent RNA-sequencing cohorts, including treatment-naïve pediatric ileal biopsies, FAO genes (*CPT1A, CPT2, SLC25A20*) were selectively downregulated in patients with or destined to develop fibrostenotic disease. Single-cell RNA-sequencing localized these transcriptional alterations specifically to fibroblasts within strictured ileum. Primary fibroblasts derived from fibrostenotic CD exhibited increased neutral lipid accumulation, impaired mitochondrial fatty acid trafficking, and diminished responsiveness to PPARγ-mediated suppression of TGFβ-induced myofibroblast activation. Together, these findings demonstrate that FAO impairment is a conserved, fibroblast-specific metabolic program associated with intestinal fibrosis in CD and suggest that metabolic modulation of stromal cells represents a potential therapeutic strategy for fibrostenotic disease.

## Introduction

Crohn’s disease (CD) is a chronic, relapsing inflammatory disorder marked by transmural inflammation affecting any portion of the alimentary tract. Clinically, CD is heterogeneous, with distinct phenotypes including inflammatory, penetrating, and stricturing presentations. Among these, stricturing CD poses a particularly challenging clinical course, affecting up to 70% of patients within a decade of diagnosis and often leading to recurrent bowel obstructions and surgical resections(1, 2). Despite its prevalence, the pathogenesis of intestinal fibrosis and stricture formation remains incompletely understood and likely reflects contributions from multiple tissue compartments(3). Moreover, no approved anti-fibrotic therapies currently exist.

Fibrosis, a hallmark of chronic tissue injury and dysregulated repair, involves the excessive accumulation of extracellular matrix (ECM) components, principally mediated by activated mesenchymal cells such as myofibroblasts (4, 5). These cells arise from diverse sources and are metabolically reprogrammed to support ECM production, contractility, and survival (6). In well-established models of pulmonary and dermal fibrosis, including the bleomycin-induced lung injury model and stiff skin syndromes, myofibroblast activation is tightly linked to shifts in cellular metabolism, notably enhanced glycolysis and suppressed fatty acid oxidation (FAO)(7–11). For instance, in idiopathic pulmonary fibrosis, inhibition of the key FAO rate limiting enzyme carnitine palmitoyltransferase 1A (CPT1A) exacerbates fibrosis, whereas enhancement of FAO ameliorates disease severity (12, 13). These metabolic adaptations are not merely consequences but drivers of fibrogenesis, offering compelling targets for therapeutic modulation.

While the fibrotic phenotype of CD shares histological and cellular features of fibrosis in other organ systems, the metabolic regulation of myofibroblast activation and fibrosis in CD remains poorly understood. In this study, we employed a multi-omic approach to uncover evidence of dysregulated fatty acid oxidation (FAO) in fibrotic CD. These findings support a model in which altered cellular metabolism contributes to myofibroblast activation. Targeting these metabolic vulnerabilities may offer new therapeutic strategies to mitigate fibrotic complications in CD.

## Results

### Cohort Demographics

The metabolomic study cohort included 59 patients, comprising 38 individuals with Crohn’s disease (CD) and 21 non–inflammatory bowel disease (NIBD) controls (table 1). CD patients were younger than NIBD controls (43 ± 15 vs 59 ± 16 years, P = 0.0002), while sex and smoking status were similar between groups. Nearly half of CD patients exhibited stricturing disease behavior. Detailed clinical characteristics are provided in Supplementary Table 1.

**Table 1.** Clinical and Demographic Characteristics of the Metabolomic Study Cohort. Demographic, clinical, and treatment characteristics of non–inflammatory bowel disease (NIBD, *n* = 21) and Crohn’s disease (CD, *n* = 38) patients included in metabolomic profiling. Montreal classification is used to describe CD disease extent, including age at diagnosis (A1: <17 years; A2: 17–40 years; A3: >40 years), behavior (B1: non-stricturing/non-penetrating; B2: stricturing; B3: penetrating; B2B3: mixed phenotype), and location (L1: ileal; L2: colonic; L3: ileocolonic). Categorical variables were compared using Fisher’s exact test; continuous variables were compared using unpaired 2-tailed Student’s *t* test. *P* <.05 was considered statistically significant. 5-aminosalicylic acid (5-ASA); TNF, anti–tumor necrosis factor.

|  | <u>NIBD</u> | <u>CD</u> | <u>P value</u> |
| --- | --- | --- | --- |
| **Total Number** | 21 | 38 | NA |
| **Sex, female/male, n (%)** | 13 (62), 8 (38) | 24 (63), 14 (37) | >0.999 |
| **Age at sampling, y, mean (SD)** | 59 (16) | 43 (15) | 0.0002 |
| **Current/former smoker, n (%)** | 10 (48) | 21 (55) | 0.598 |
| **Disease Extent (Montreal Classification)** |  |  |  |
| Age at diagnosis (A1 / A2 / A3), n (%) | NA | 8 (21) / 25 (66) / 5 (13) | NA |
| Behavior (B1 / B2 / B3 / B2B3), n (%) | NA | 7 (18) / 18 (47) / 3 (8) / 10 (26) | NA |
| Location (L1/L2/L3), n (%) | NA | 14 (37) / 4 (11) / 20 (53) | NA |
| **Treatment history** |  |  |  |
| Steroids, n (%) | 2 (10) | 35 (92) | < 0.0001 |
| 5-ASA, n (%) | 0 (0) | 23 (61) | < 0.0001 |
| Immunomodulators, n (%) | 0 (0) | 33 (87) | < 0.0001 |
| Anti-TNF, n (%) | 0 (0) | 30 (79) | < 0.0001 |
| Non-anti-TNF biologic, n (%) | 0 (0) | 8 (21) | < 0.0001 |
| **Other Diagnoses** |  |  |  |
| Hypertension, n (%) | 11 (52) | 9 (24) | 0.0433 |
| Hyperlipidemia, n (%) | 6 (29) | 5 (13) | 0.175 |
| Diabetes, n (%) | 3 (14) | 2 (5) | 0.337 |

### Global Metabolomic and Lipidomic Profiling Identifies Neutral Lipid and Acylcarnitine Accumulation in Crohn’s Disease

Untargeted metabolomic and lipidomic profiling was performed on surgically resected, non-inflamed colonic tissue (confirmed on histology) obtained from CD and NIBD adult patients. Comprehensive analysis was conducted using two high-dimensional platforms: global metabolomics (HD4 dataset) and complex lipidomics (CLP dataset).

Multivariate modeling with sparse partial least squares discriminant analysis (sPLS-DA) demonstrated clear separation between CD and NIBD samples in both global metabolomic and lipidomic datasets (Figure 1A-B), indicating distinct metabolic signatures associated with Crohn’s disease. To further identify the metabolites driving this separation, variable correlation loading plots were generated (Figure 1C-D). In the global metabolomics dataset, discriminating features included several amino acids, peptides, and pathway-associated metabolites. Within the lipidomics dataset, triacylglycerols (TAGs) emerged as key drivers of group separation, with additional contributions from diacylglycerols (DAGs), phosphatidylcholines (PCs), and hexosylceramides (HCERs).

**Figure 1.**
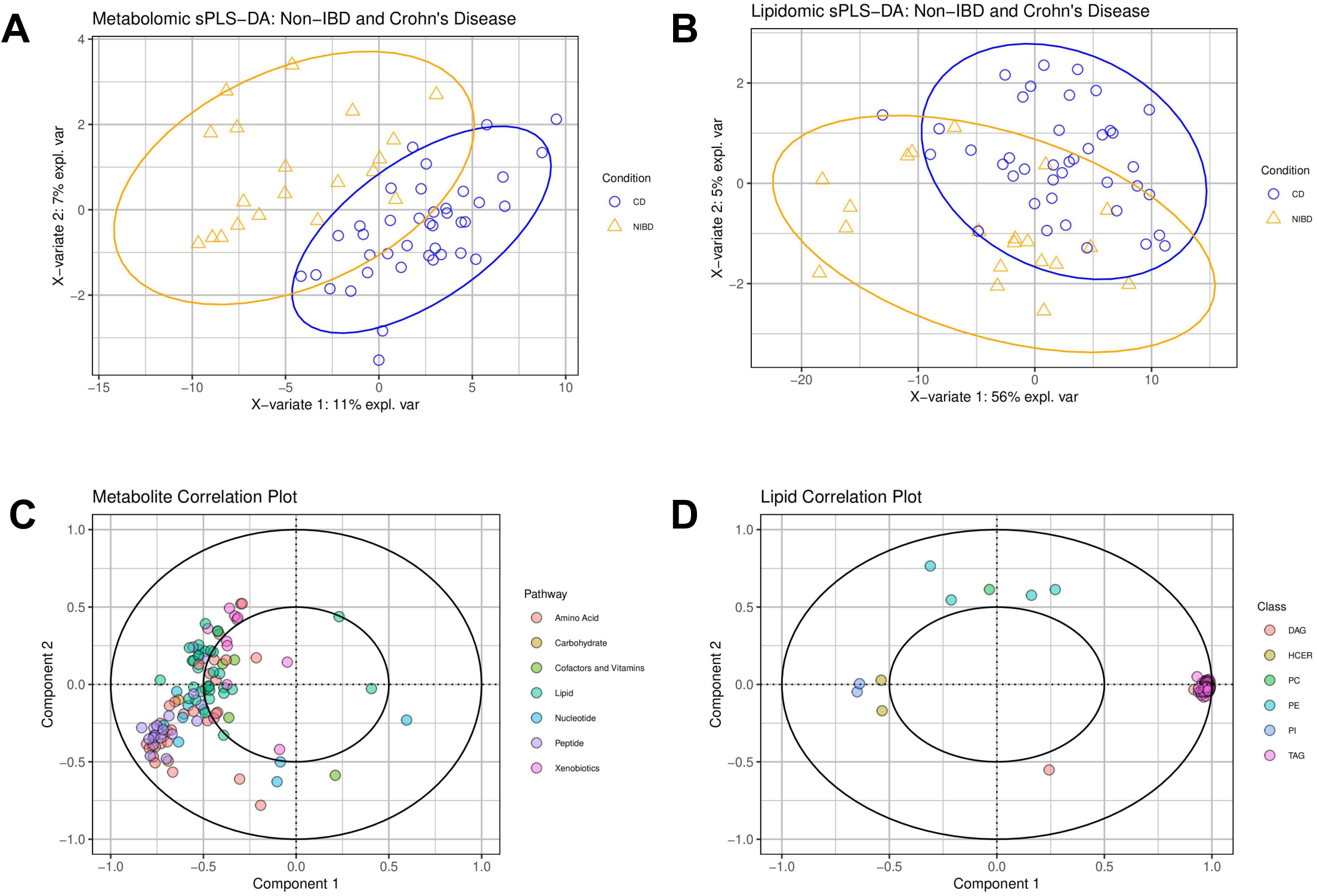
Global metabolomic and lipidomic profiling of non-inflamed colonic tissue from Crohn’s disease (CD) patients and non-IBD (NIBD) controls. (A) Sparse partial least squares discriminant analysis (sPLS-DA) of global metabolomics (HD4 dataset) demonstrates clear separation between CD (blue) and NIBD (orange) samples, indicating disease-associated metabolic reprogramming independent of active mucosal inflammation. Each point represents one patient sample; ellipses denote 95% confidence intervals. (B) sPLS-DA of the complex lipidomics (CLP) dataset similarly reveals distinct separation between CD and NIBD samples, confirming a disease-specific lipidomic signature in non-inflamed colonic tissue. (C) Variable correlation loading plot corresponding to the HD4 metabolomics model, identifying amino acids, peptides, and pathway-associated metabolites as key contributors to group discrimination. Variables positioned further from the origin exert greater influence on separation. (D) Variable correlation loading plot corresponding to the CLP lipidomics model, identifying triacylglycerols (TAGs) as the predominant drivers of CD versus NIBD separation, with additional contributions from diacylglycerols (DAGs), phosphatidylcholines (PCs), and hexosylceramides (HCERs). Abbreviations: DAG, diacylglycerol; HCER, hexosylceramide; PC, phosphatidylcholine; PE, phosphatidylethanolamine; PI, phosphatidylinositol; TAG, triacylglycerol.

Given the lipidomic separation observed by sPLS-DA, we next performed lipid subclass analyses to characterize specific lipid species contributing to the observed group differences. Consistent with the variable loading plots from sPLS-DA (Figure 1D), which identified multiple TAG species as key drivers of group separation, heatmap visualization of TAG subclasses stratified by fatty acid chain length and degree of unsaturation demonstrated a broad and consistent elevation of TAGs in Crohn’s disease compared to NIBD controls (Figure 2A).

**Figure 2.**
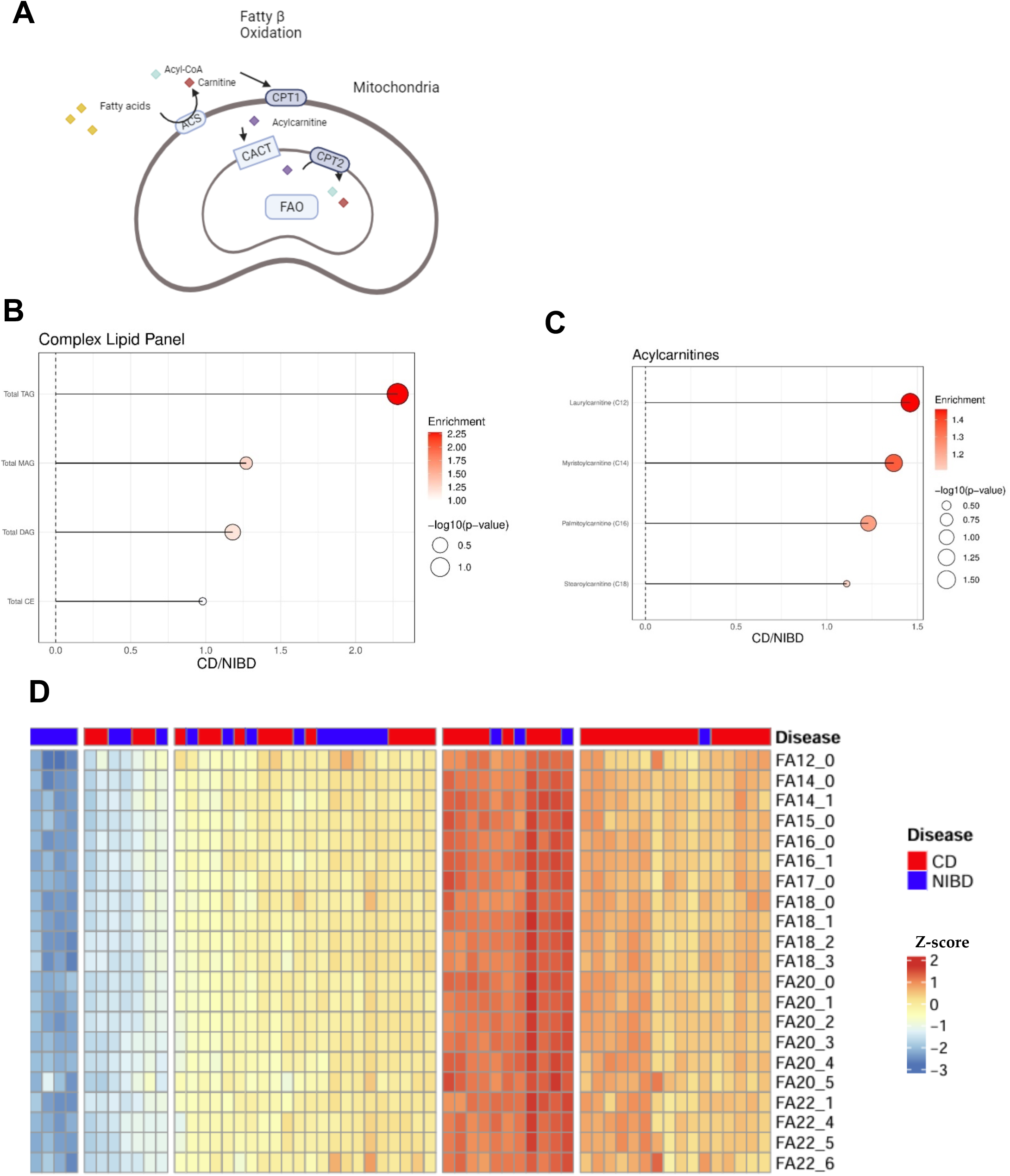
Triacylglycerols and acylcarnitines are enriched in Crohn’s disease. (A) Schematic representation of the mitochondrial fatty acid oxidation (FAO) pathway, illustrating key enzymes involved in fatty acid transport and oxidation, including CPT1, CACT, and CPT2, which were assessed in downstream transcriptomic analyses. *Enzyme abbreviations: CPT1, carnitine palmitoyltransferase 1; CACT, carnitine-acylcarnitine translocase; CPT2, carnitine palmitoyltransferase 2*. (B) Lollipop plot of complex lipid subclasses showing fold-change (CD/NIBD) and statistical significance for total triacylglycerol (TAG), monoacylglycerol (MAG), diacylglycerol (DAG), and cholesteryl ester (CE) levels. Total TAG concentrations were significantly elevated in CD compared to NIBD (fold-change 2.28; *P* = 0.0387), while MAG, DAG, and CE levels were not significantly different between groups. (C) Lollipop plot of selected acylcarnitine species showing fold-change enrichment in CD versus NIBD controls. Laurylcarnitine (C12) (1.46-fold; *P* = 0.0254) and myristoylcarnitine (C14) (1.37-fold; *P* = 0.0416) were significantly increased, while palmitoylcarnitine (C16) showed a non-significant trend (1.23-fold; *P* = 0.0783) and stearoylcarnitine (C18) was not significantly different (*P* = 0.3579). (D) Heatmap of triacylglycerol (TAG) species comparing Crohn’s disease (CD) and non-inflammatory bowel disease (NIBD) control samples. TAGs are grouped by total fatty acid carbon chain length and number of double bonds. Relative abundances are Z-score normalized.

To further quantify lipid class differences, we analyzed summed concentrations of key neutral lipids and fatty acid oxidation intermediates (Figure 2B-C). Total TAG concentrations were significantly elevated in CD relative to controls (*P*=0.0387), whereas levels of DAGs, monoacylglycerols (MAGs), and cholesteryl esters (CEs) did not significantly differ between groups.

In addition to TAG accumulation, multiple acylcarnitine species demonstrated selective enrichment in CD samples compared to controls. Laurylcarnitine (C12 acylcarnitine) was significantly increased in CD patients (fold-change 1.46; *P* = 0.0254), as was myristoylcarnitine (C14 acylcarnitine) (fold-change 1.37; *P* = 0.0416). Palmitoylcarnitine (C16 acylcarnitine) showed a non-significant trend toward elevation (fold-change 1.23; *P* = 0.0783), while stearoylcarnitine (C18 acylcarnitine) levels were not significantly different (*P* = 0.3579). The parallel increase in both neutral lipid storage species and selected acylcarnitines suggests a potential impairment in mitochondrial fatty acid oxidation (Figure 2D) in Crohn’s disease tissue.

### Coordinated Suppression of FAO–PPARγ Signaling with Reciprocal Induction of Glycolytic and Fibrogenic Programs in Crohn’s Disease

To relate the lipid level signature in colon tissue to underlying transcriptional programs implicated in lipid metabolism, we next examined a curated panel of genes representing fatty acid oxidation (FAO), Peroxisome Proliferator Activated Receptor Gamma (PPARγ) signaling, glycolysis, and Transforming Growth Factor Beta (TGF β) pathways in three independent RNA sequencing cohorts: adult non inflamed colon, adult inflamed ileum, and non-inflamed treatment naïve pediatric ileum (Figure 3A–C). For each dataset, we calculated DESeq2 log2 fold change values for Crohn disease groups relative to non IBD controls and summarized them in pathway annotated heatmaps.

**Figure 3.**
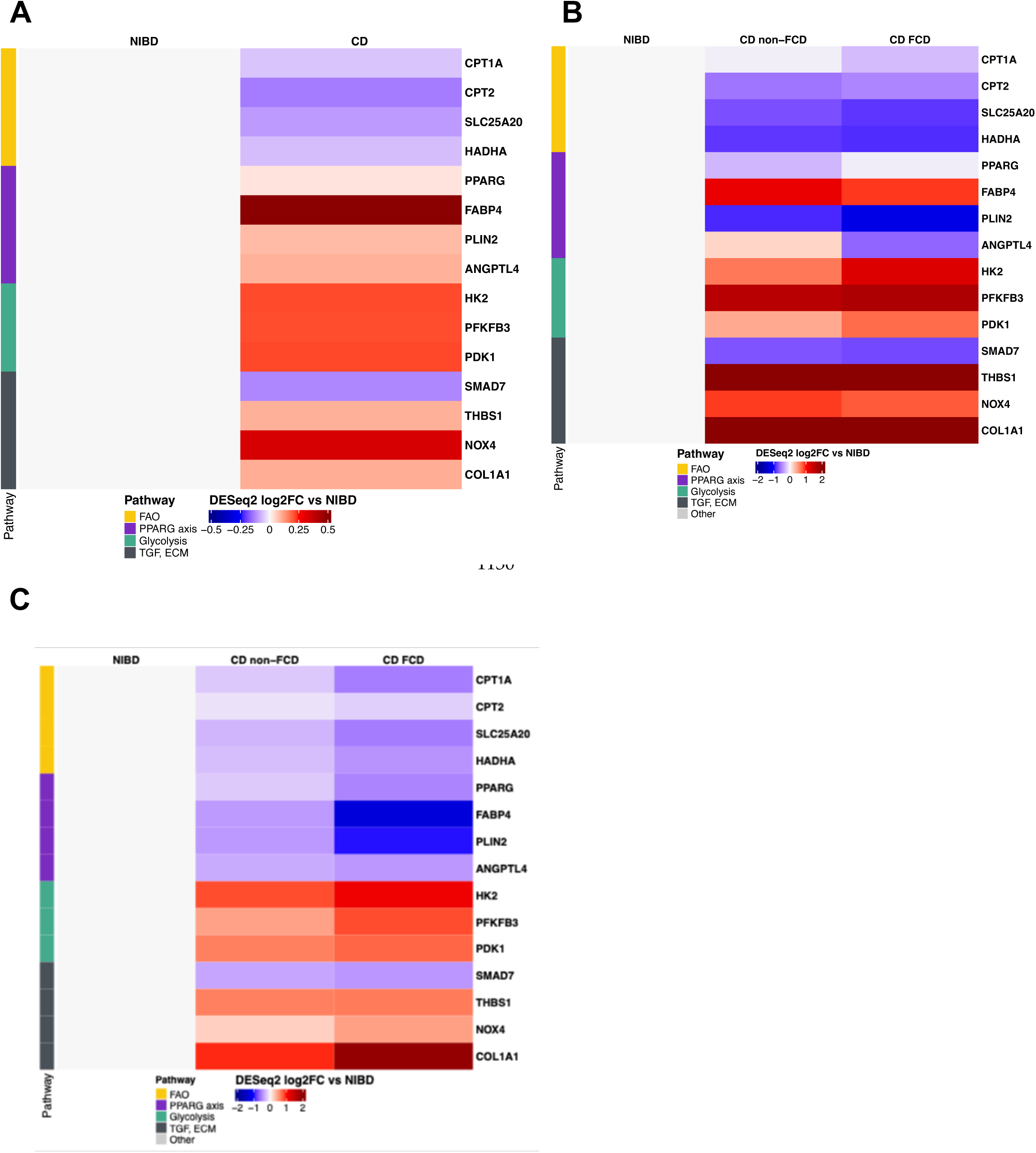
Coordinated transcriptional remodeling of fatty acid oxidation, glycolysis, PPARγ, and TGF β programs across adult colon, adult ileum, and pediatric ileum in Crohn disease. (A) Heatmap of DESeq2 log2 fold change (log2FC) for selected fatty acid oxidation (CPT1A, CPT2, SLC25A20, HADHA), PPARγ axis (PPARG, FABP4, PLIN2, ANGPTL4), glycolytic (HK2, PFKFB3, PDK1), and TGF and extracellular matrix related genes (SMAD7, THBS1, NOX4, COL1A1) in non-inflamed adult colonic tissue, comparing Crohn disease (CD) to non-inflammatory bowel disease (NIBD) controls. Colors indicate direction and magnitude of differential expression (blue, decreased; red, increased; scale −0.5 to +0.5 log2FC relative to NIBD). (B) Corresponding heatmap from adult ileal tissue, showing transcriptional changes in non-inflamed NIBD, inflamed CD patients with fibrostenotic Crohn disease (CD FCD) at the time of tissue collection, and inflamed CD patients without fibrostenosis (CD non FCD), each displayed as log2FC relative to NIBD. Colors indicate direction and magnitude of differential expression (blue, decreased; red, increased; scale −2 to +2 log2FC relative to NIBD). (C) Analogous analysis in treatment naive pediatric ileal biopsies obtained at diagnosis from NIBD, non FCD, and patients who were classified as FCD based on development of fibrostenotic disease during longitudinal follow up, showing that many FCD associated transcriptional changes are already present at the time of diagnosis, before strictures are clinically apparent. Colors indicate direction and magnitude of differential expression (blue, decreased; red, increased; scale −2 to +2 log2FC relative to NIBD).

In the adult colonic cohort, Crohn disease samples showed a coherent shift toward reduced FAO and PPARγ axis signaling with reciprocal induction of glycolytic and fibrogenic genes (figure 3A). FAO components Carnitine palmitoyltransferase 1A (*CPT1A*), Carnitine palmitoyltransferase 2 (*CPT2*), Solute carrier family 25 member 20 (also known as the carnitine-acylcarnitine translocase, (*SLC25A20* or *CACT*) and Hydroxyacyl-CoA dehydrogenase trifunctional multienzyme complex subunit alpha (*HADHA*) tended to be lower in CD colon compared with non IBD, whereas glycolytic regulators Hexokinase 2 (*HK2*) and 6-phosphofructo-2-kinase/fructose-2,6-bisphosphatase 3 (*PFKFB3*) and the TGF/ECM effectors Thrombospondin 1 (*THBS1*), NADPH Oxidase 4 (*NOX4)* and Collagen Type 1 alpha chain (*COL1A1*) were generally higher. PPARγ pathway members *PPARG*, Perilipin 2 (*PLIN2*) and Fatty acid binding protein 4 (*FABP4*) were reduced, consistent with diminished lipid handling capacity. While the log fold values were modest compared to the rest of the cohort, it still highlighted the overall direction of change supported a coordinated down regulation of FAO and PPARγ signaling in adult colon.

We observed a similar but more stratified pattern in the adult ileal resection cohort, where patients were grouped at the time of surgery as non stricturing CD (CD non-FCD) or fibrostenotic CD (CD FCD) based on ileal stricture status at the time of sample collection (Figure 3B). Compared with non IBD ileum, FAO genes showed the greatest reduction in CD FCD, with CD non-FCD generally intermediate. Glycolytic and TGF/ECM genes were most strongly induced in CD FCD, again with a graded pattern in CD non-FCD. For example, HK2 and PFKFB3, as well as THBS1 and COL1A1, were more highly up regulated in CD FCD than in CD non-FCD, mirroring the more advanced fibrotic phenotype. As in the colon, several genes such as SMAD7 showed less consistent modulation, suggesting some pathway elements are more tightly coupled to fibrostenotic status than others.

Adult tissues represented patients with an already manifest clinical phenotype. To understand potential prognostic value of altered FAO programs in FCD we then asked whether a similar transcriptional program is already evident at diagnosis in children who later develop stricturing disease. In this cohort, all ileal biopsies were obtained at treatment naïve diagnosis before the patient had developed a stricture, and individuals were prospectively classified as non-FCD or FCD based on longitudinal follow up (Figure 3C)(14). Despite being sampled prior to stricture formation, children who later progressed to FCD already showed marked down regulation of FAO genes *CPT1A, CPT2* and *SLC25A20* relative to non IBD, with non FCD patients again displaying more modest or absent changes. PPARγ axis genes *PPARG, PLIN2* and *FABP4* were similarly reduced in future FCD, whereas *PLIN2* tended to be decreased across both CD groups. In contrast, glycolytic genes *HK2* and *PFKFB3* and the profibrotic TGF/ECM mediators *THBS1, NOX4* and *COL1A1* were preferentially up regulated in the future FCD group. As in adults, *SMAD7* did not decrease uniformly across all comparisons, indicating that not every TGF regulatory node tracks strictly with fibrosis risk.

Collectively, these data demonstrate a conserved transcriptional reprogramming across adult and pediatric Crohn’s disease characterized by suppression of the FAO–PPARγ axis and reciprocal activation of glycolytic and fibrogenic pathways, with early emergence in patients destined for fibrostenotic disease.

### Single-Cell RNA-Sequencing Identifies Compartment-Specific Decreases in FAO Gene Expression in Fibrotic Ileum

Because stromal cells, including fibroblasts and smooth muscle cells, are primary mediators of intestinal fibrosis, we next sought to evaluate whether the decreases in FAO gene expression observed in bulk RNA-sequencing were reflected within specific stromal compartments. To address this, we re-interrogated single-cell RNA-sequencing data derived from patients with Crohn’s disease (15). This dataset comprises tissue from 20 human gut resections, including 7 normal controls and 13 Crohn’s disease samples. The CD samples were anatomically stratified into non-involved, inflamed, and stenotic regions, enabling compartment-specific analysis of transcriptional changes across the progression of intestinal fibrosis.

Given the pronounced suppression of FAO genes observed in bulk RNA-sequencing and the enrichment of fibroblast activation markers in fibrostenotic tissue, we focused our single-cell analysis on fibroblast populations to determine whether these transcriptional changes were cell type–specific. Representative fibroblast-associated genes spanning key metabolic and fibrogenic pathways are shown in Figure 4A. Consistent with a profibrotic phenotype, *COL1A1* expression demonstrated a stepwise increase from normal through strictured ileum, confirming fibroblast activation within fibrotic tissue. Expression of the fatty acid oxidation (FAO) enzyme *HADHA* (required for catalyzing final steps of and the PPARγ target gene *FABP4* decreased in strictured fibroblasts, aligning with impaired FAO and reduced PPARγ signaling observed in bulk tissue and primary fibroblast assays. In contrast, glycolytic and inflammatory regulators, including *HIF1A, IER3*, and *IL6*, were upregulated in inflamed and strictured regions. *TGFB3* expression also increased with disease progression, further supporting enhanced TGFβ pathway activity in fibroblasts from fibrotic ileum.

**Figure 4.**
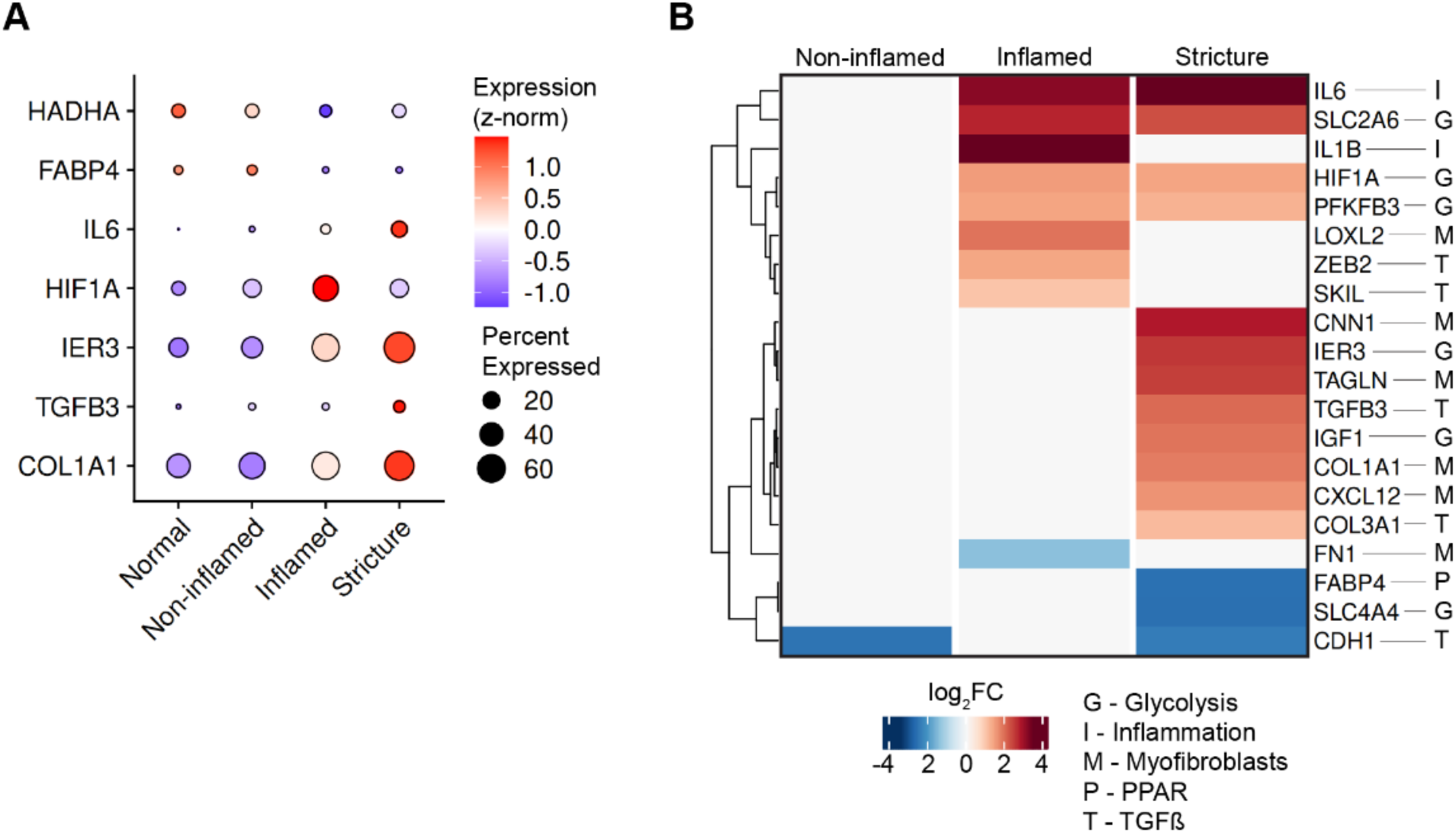
Single-cell RNA-sequencing reveals fibroblast-specific metabolic reprogramming and TGF-β pathway activation in fibrotic Crohn’s disease ileum. (A) Quantitative dot plot showing representative fibroblast genes across key metabolic and fibrogenic pathways, including glycolysis (*IER3, HIF1A*), PPARγ signaling (*FABP4*), fatty acid oxidation (*HADHA*), inflammation (*IL6*), myofibroblast activation (*COL1A1*), and TGF-β signaling (*TGFB3*). Color values denote z-normalized gene expression, and dot size represents the proportion of fibroblasts expressing each transcript across normal, non-inflamed, inflamed, and strictured ileal tissue. (B) Pseudobulk differential expression heatmaps showing log₂ fold-change (log₂FC) values for fibroblast-expressed genes relative to normal controls. Data represents aggregated fibroblast transcriptomes from the ECLP2 dataset. Only genes and contrasts with significant changes (adjusted *P* ≤ 0.1) are shown. Dendrograms indicate hierarchical clustering across gene categories.

To complement these single-gene trends, pseudobulk differential expression analysis was performed using aggregated fibroblast transcriptomes across tissue states (Figure 4B).

Fibroblasts from strictured tissue showed coordinated upregulation of Genes exhibiting significant differences (adjusted *P* ≤ 0.1) were annotated by pathway category. Fibroblasts from strictured tissue showed coordinated upregulation of glycolytic and TGFβ-associated transcripts (*IER3, HIF1A, PFKFB3, TGFB3*), along together with myofibroblast and extracellular matrix genes (*COL1A1, COL3A1, FN1, CNN1, TAGLN*). In contrast, while FAO-and PPARγ-related genes (*FABP4, HADHA*) were significantly downregulated. These results identify a fibroblast-specific transcriptional program in strictured ileum characterized by suppression of FAO and PPARγ signaling and concomitant activation of glycolytic and TGFβ pathways, consistent with metabolic reprogramming toward a fibrotic myofibroblast phenotype.

Together, these single-cell findings localize the metabolic alterations identified in bulk RNA-sequencing to the stromal compartment, specifically fibroblasts, and reinforce that impaired FAO and PPARγ signaling accompany transcriptional activation of fibrogenic and glycolytic pathways during fibroblast differentiation in Crohn’s disease.

### Myofibroblasts from Fibrotic Crohn’s Disease Exhibit Neutral Lipid Accumulation

Given the observed downregulation of FAO-related genes in fibroblast populations identified by single-cell RNA sequencing (Figure 4), we next sought to interrogated FAO in primary fibroblasts isolated from the terminal ileum (, the primary site of fibrostenotic disease) from, in 3 non-fibrotic CD (non-FCD patients), and 4 FCD patients, and in 4 NIBD controls. Isolated cells were cultured and passaged under standard conditions; flow cytometry for CD90 or αSMA were used to assess homogeneity of isolated fibroblasts and myofibroblast differentiation respectively (Supplementary Figure 1).

We examined lipid droplet (LD) accumulation in these purified primary myofibroblasts to validate metabolomic evidence of TAG accumulation in FCD tissues. Confocal microscopy using the neutral lipid dye BODIPY 493/503 revealed visibly increased cytoplasmic LD content in fixed FCD-derived myofibroblasts compared to those from non-FCD and NIBD controls (representative figure 5A). Quantification of LD density showed increased accumulation in FCD samples relative to non-FCD and NIBD samples (Figure 5B), consistent with a shift toward intracellular neutral lipid storage. We also measured BODIPY signal intensity by flow cytometry in live, cultured intestinal myofibroblasts; a representative staining profile is shown in Figure 5C, and the gating strategy is presented in Supplementary Figure 2. Fluorescence intensity of lipid drops was higher in FCD samples compared to non-FCD and NIBD controls, further supporting enhanced neutral lipid accumulation in fibrotic disease (Figure 5D).

**Figure 5.**
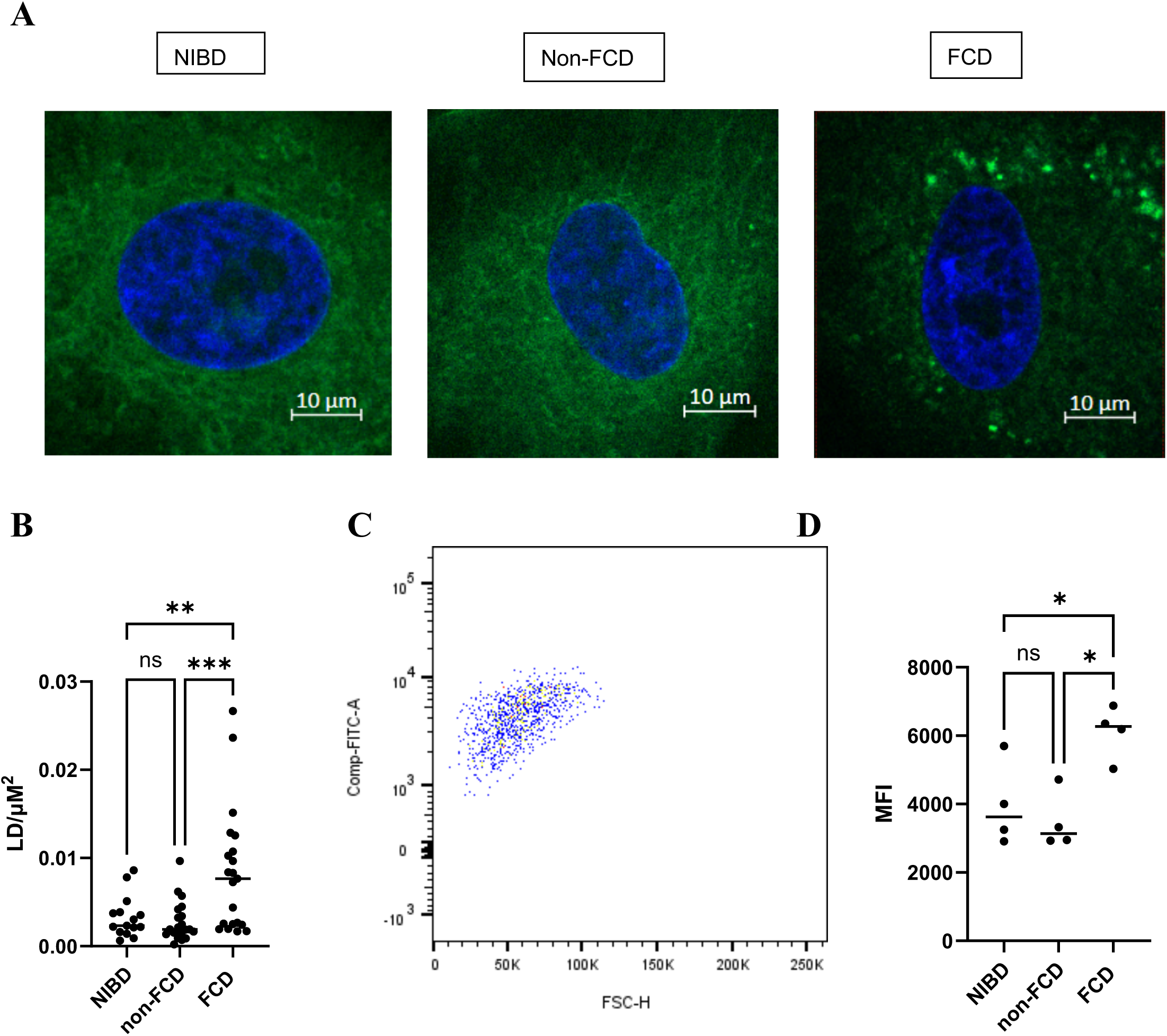
Myofibroblasts from fibrotic Crohn’s disease exhibit neutral lipid accumulation. (A) Representative confocal images of BODIPY 493/503–stained intestinal myofibroblasts isolated from terminal ileum of NIBD, non-FCD, and FCD patients. Nuclei are labeled with Hoechst 33342. (B) Lipid droplet density quantified as lipid droplets per cytoplasmic area in cultured myofibroblasts. (C) Representative flow cytometry profile of BODIPY staining in live myofibroblasts. (D) Mean fluorescence intensity (MFI) of BODIPY 493/503 in NIBD, non-FCD, and FCD myofibroblast cultures. Quantification includes pooled cells from NIBD, non-FCD, and FCD samples. Flow cytometry was performed on n = 4 NIBD, n = 4 non-FCD, and n = 4 FCD patient-derived myofibroblast lines. Gating strategy shown in Supplementary Figure Y. Statistical analysis by one-way ANOVA. *P < 0.05, **P < 0.01, ***P < 0.001.

These results suggest that intestinal myofibroblasts from fibrotic CD tissue harbor increased neutral lipid content, supporting a model of impaired lipid utilization and fatty acid oxidation defects in fibrosis-associated stromal cells.

### Impaired Mitochondrial Fatty Acid Trafficking in Fibrotic Crohn’s Disease

We sought to spatiotemporally resolve neutral lipid accumulation by examining fatty acid trafficking into mitochondria of intestinal myofibroblasts. Myofibroblasts were pulsed with BODIPY-C16, then transferred to fresh, unlabeled media to allow for intracellular redistribution (Figure 6A). Mitochondria and nuclei were respectively counterstained, and cells were visualized by confocal imaging (representative staining in Figure 6B).

**Figure 6.**
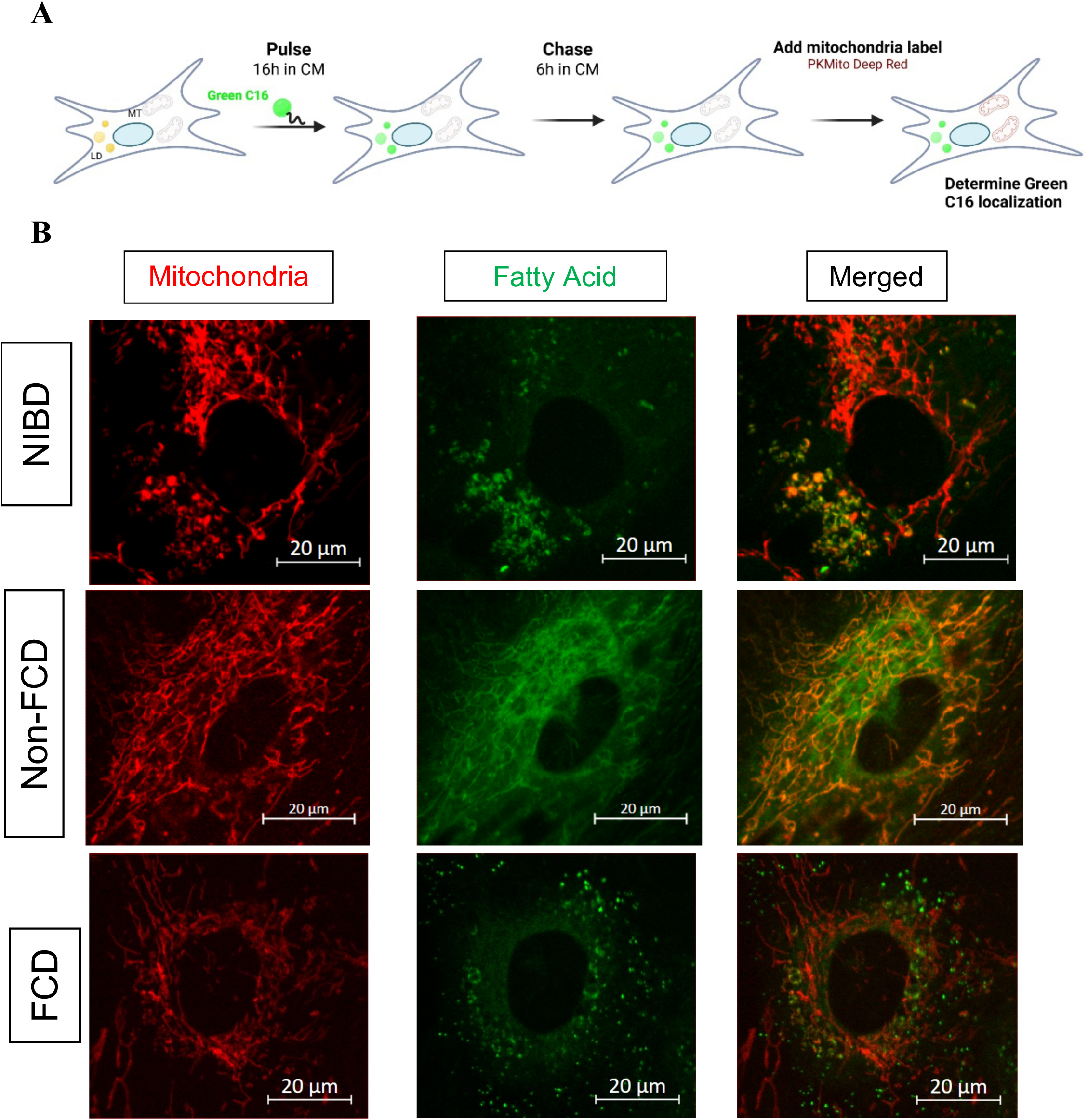
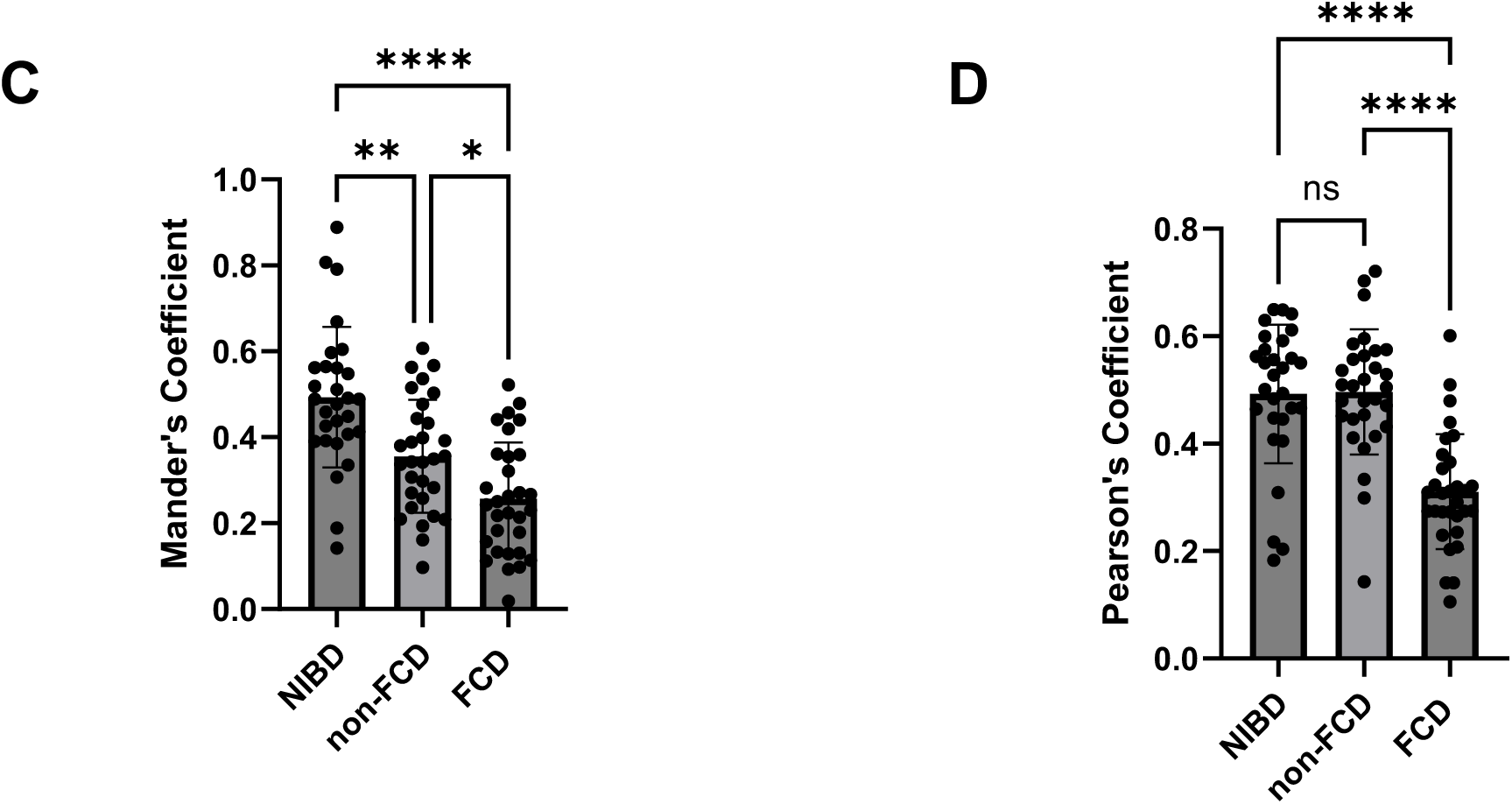
Fibrotic Crohn’s disease (FCD) myofibroblasts exhibit impaired mitochondrial fatty acid trafficking. **(A)** Schematic of the pulse-chase experiment. Myofibroblasts were incubated with BODIPY-C16 to label intracellular fatty acid pools, followed by a chase in fresh media and subsequent staining with PKmito Deep Red to visualize mitochondria. **(B)** Representative confocal image of BODIPY-C16 (green) and PKmito Deep Red (magenta) colocalization in myofibroblasts. **(C)** Quantification of colocalization using Manders’ M1 coefficient revealed significantly reduced overlap of mitochondrial and fatty acid signals in non-FCD and FCD compared to NIBD myofibroblasts (P < 0.0011 for NIBD vs. non-FCD; P < 0.0001 for NIBD vs. FCD; P = 0.0219 for non-FCD vs. FCD). **(D)** Pearson’s correlation coefficient was also significantly reduced in FCD compared to both NIBD and non-FCD myofibroblasts (P < 0.0001 and P = 0.0001), while NIBD and non-FCD were not significantly different. Pooled cells for, NIBD, non-FCD, and FCD samples were analyzed. Statistical analysis was performed using one-way ANOVA with Tukey’s multiple comparisons test for Manders’ coefficient, and Kruskal–Wallis test with Dunn’s post-test for Pearson’s correlation coefficient.

The degree of colocalization (fraction of mitochondria label overlapping with BODIPY-C16 FA signal) was calculated using Manders’ M1 coefficient and Pearson’s correlation coefficient, which reflects the extent to which mitochondria and BODIPY-C16 FA intensities align on a pixel-by-pixel basis. Both measures demonstrated progressive reductions in colocalization from NIBD to non-FCD to FCD myofibroblasts. Manders’ coefficient was significantly lower in both non-FCD and FCD compared to NIBD myofibroblasts (Figure 6C), suggesting impaired FFA utilization in mitochondria. Similarly, Pearson’s correlation coefficient was significantly reduced in FCD relative to both NIBD and non-FCD (Figure 6D), indicating decreased spatial correlation of mitochondrial and FA signals.

These findings suggest that mitochondrial fatty acid trafficking is impaired in fibrotic myofibroblasts, aligning with prior observations of altered lipid accumulation and reduced FAO gene expression.

### Fibrotic Myofibroblasts Exhibit Resistance to PPARγ-Mediated Suppression of Activation

Given our transcriptional evidence of impaired FAO and reduced PPARγ activity in intestinal fibroblasts from Crohn’s disease, we next examined whether these molecular alterations influence fibroblast responsiveness to PPARγ agonism during myofibroblast activation. Rosiglitazone, a selective PPARγ agonist known to promote fatty acid oxidation and suppress profibrotic fibroblast activation, was therefore used to test whether enhancing PPARγ signaling modulates TGFβ-induced myofibroblast activation. Using CCD-18CO fibroblast cell lines and NIBD primary fibroblasts, we first established that rosiglitazone effects are due to modulation of PPARγ and not impaired cell viability (Supplementary Figure 3). Next, primary fibroblasts from NIBD controls (N=2), and fibrotic and non-fibrotic CD patients (representing 3 independent patients) were stimulated with rosiglitazone (10 μM) and concomitant TGFβ (10 ng/mL) to induce αSMA activation which was assessed using western blotting; representative blots are shown in Figures 7A, B respectively for NIBD and non-FCD. (Figure 7A-B). Densitometry analysis indicates that only a modest αSMA induction was observed in NIBD-derived ileal fibroblasts (Figure 7C), which was reverted to baseline with rosiglitazone treatment. In contrast, CD-derived ileal fibroblasts (Figure 7D) robustly increase TGFβ-dependent αSMA expression; rosiglitazone exposure reduced this upregulation, but to a level higher than at baseline.

**Figure 7.**
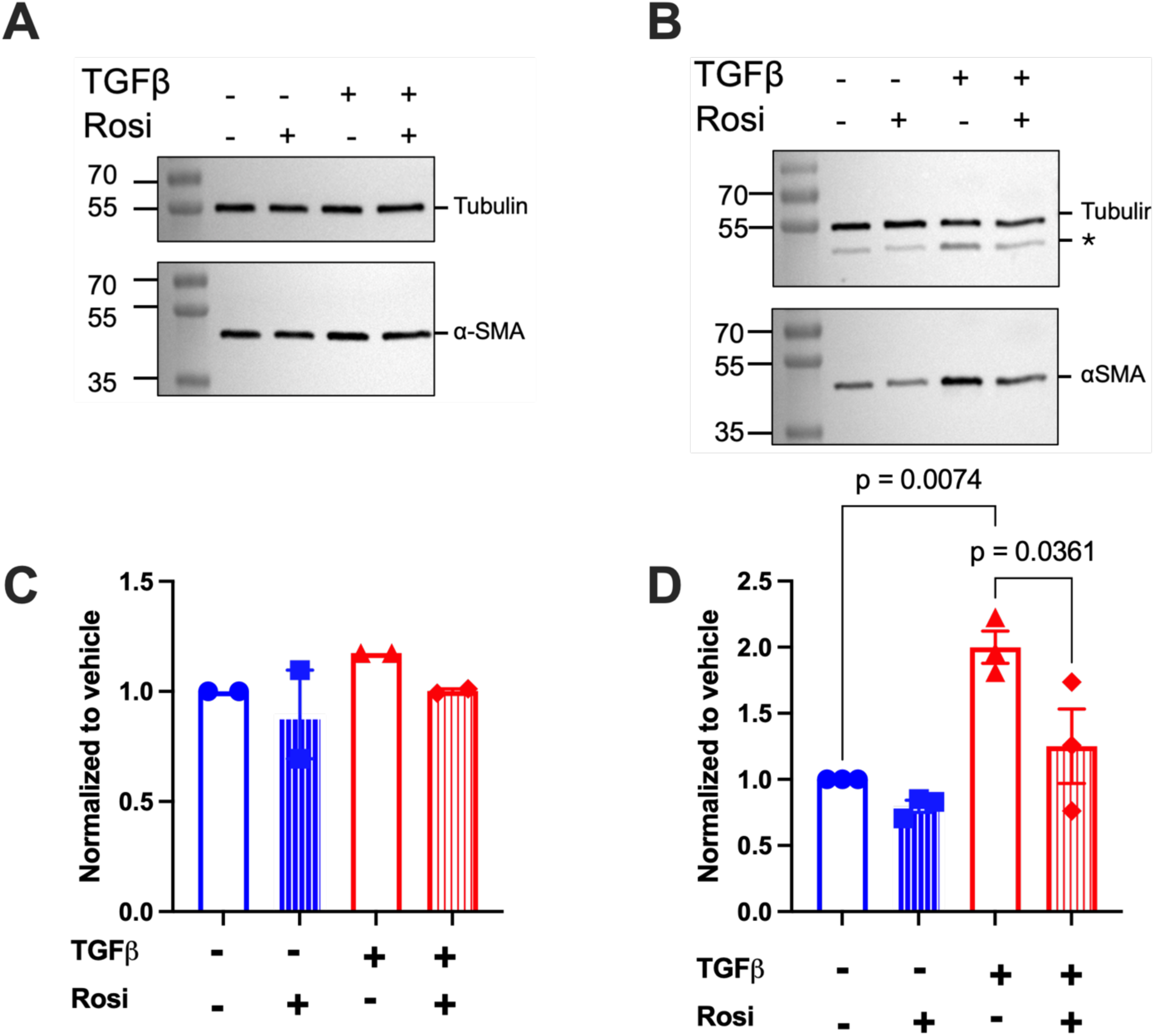
Differential responses of NIBD and CD ileal myofibroblasts to TGFβ stimulation and PPARγ agonism. (A, **B)** Representative Western blot showing ACTA2 (αSMA) expression in primary myofibroblasts isolated from the terminal ileum of (**A**) NIBD and (**B**) non-FCD patients. Cells were stimulated with TGFβ (10 ng/mL), rosiglitazone (10 μM), or both in combination. Tubulin was used as a loading control. * in panel B indicates αSMA residual signal following blot stripping. (**C, D**) Densitometric quantification of αSMA expression normalized to Tubulin. Compared to NIBD-derived ileal cells, CD ileal myofibroblasts showed significantly stronger induction of αSMA with TGFβ. Statistical analysis was performed using one-way ANOVA with Sidak’s post-hoc correction.

These findings indicate that fibroblasts from Crohn’s disease exhibit diminished sensitivity to PPARγ activation during TGFβ-induced myofibroblast differentiation, consistent with transcriptional signatures of reduced FAO and reduced PPARγ pathway activity identified in both bulk and single-cell RNA sequencing.

## Discussion

Crohn’s disease (CD) is a chronic relapsing inflammatory disorder of the gastrointestinal tract with rising global incidence and disease burden (16, 17). Despite the advent of biologic therapies, including anti-TNF agents, anti-integrins, IL-12/23 inhibitors, and more recently JAK inhibitors, response rates remain variable, and a substantial proportion of patients experience treatment failure and ultimately require surgical intervention (18, 19). Failure may result from refractory inflammation, but increasingly recognized is the progression of complications independent of active inflammation, including fistulas and fibrostenotic strictures(20–23). Notably, even patients who achieve clinical or endoscopic remission may continue to develop stricturing disease, highlighting a critical limitation of current therapies(20, 21). Most available treatments target the immune component of CD, yet little is known about the non-immune mechanisms that drive tissue remodeling and fibrosis(24). Importantly, no approved therapies currently exist to prevent or reverse intestinal fibrosis(19, 22). This unmet clinical need underscores the importance of understanding the non-immune drivers of fibrotic complications. In this multi-omic study, we identify alterations in cellular metabolism as a potential mechanistic contributor to fibrostenotic disease.

To explore novel disease pathways, we employed an untargeted metabolomic approach to identify metabolic perturbations in CD. Using non-inflamed colon tissue from CD and non-IBD (NIBD) controls, we observed striking enrichment of triacylglycerols (TAGs) and acylcarnitines in CD samples. These metabolite profiles suggest disrupted lipid handling, raising the possibility of impaired fatty acid oxidation (FAO), as both TAGs and acylcarnitines serve as key precursors (25). Although dietary intake and gut microbial composition may influence tissue lipid composition, demographic and clinical features did not differ significantly between CD and NIBD controls. Dysbiosis, commonly observed in CD and known to alter microbial-derived lipid metabolites, may partially account for these differences(26). Additionally, dietary factors are also recognized modulators of host lipid profiles and influence gut microbiota (27, 28). FAO plays a central role in cellular homeostasis and is increasingly implicated in inflammation and tissue remodeling in multiple organ systems (29, 30). However, the importance of FAO in cellular phenotype prompted further transcriptomic investigation to determine whether intrinsic defects in lipid metabolism underlie CD and disease progression.

To further investigate the metabolic alterations suggested by our lipidomic data, we examined the expression of genes critical for fatty acid oxidation (FAO), including *CPT1A, CPT2*, and *CACT*, which coordinate long-chain fatty acid transport and mitochondrial oxidation(25). Bulk RNA sequencing of a matched adult colonic cohort revealed reduced expression of *CPT2* in CD samples compared to non-IBD (NIBD) controls. Given the ileum’s predilection for fibrostenotic complications, we next profiled FAO gene expression in adult ileal samples and observed a broader pattern of downregulation. Notably, *CACT* expression was reduced in both non-FCD and FCD relative to NIBD, with no significant difference between the two CD phenotypes. A similar trend was observed for *CPT2*. These findings suggest that FAO suppression may be a general feature of Crohn’s disease, but this later-stage cohort may not effectively distinguish fibrotic versus non-fibrotic phenotypes. To assess whether FAO suppression is more specific to fibrostenotic disease, we analyzed treatment-naive pediatric ileal samples stratified by phenotype. In this earlier-stage cohort, we observed selective downregulation of *CACT* and *CPT2* in FCD samples compared to both non-FCD and NIBD controls, while *CPT1A* was more strongly reduced in FCD than in non-FCD relative to NIBD. These results suggest that early FAO dysregulation may contribute to fibrostenotic pathogenesis. In parallel, we observed increased expression of glycolytic regulators (*HK1, HK2, PFKFB3, PDK1*) (6, 10), reduced expression of *PPARγ* and its downstream targets (*PLIN2, FABP4*)(31, 32), and skewing of the TGF-β pathway with decreased *SMAD7, ID1*, and *ID3* alongside increased *THBS1, SNAI1, NOX4*, and *MMP2* (33–38). These results suggest that early metabolic reprogramming involving both suppression of FAO and compensatory induction of glycolysis, coupled with loss of PPARγ restraint and enhanced TGF-β signaling may contribute to fibrostenotic pathogenesis.

To identify the cellular origin of these metabolic changes, we focused on the stromal compartment the primary site of fibrotic remodeling and leveraged single-cell RNA sequencing data from ileal resections of CD patients stratified by disease involvement.(15) As expected, *COL1A1* expression, a key driver of extracellular matrix (ECM) deposition, was elevated in fibroblasts and smooth muscle cells from strictured regions, consistent with their roles in fibrosis. FAP, another marker of activated intestinal fibroblasts, was similarly enriched in strictured fibroblasts(39). We also found that fibroblasts exhibited reduced expression of *SMAD7, ID1*, and *ID3*, along with increased *SNAI1*, further highlighting their bias toward TGF-β–driven activation. Notably, we observed decreased expression of *CPT1A* specifically in fibroblasts from strictured tissue compared to those from uninflamed and inflamed regions, along with a similar trend for *CACT*. These findings highlighted fibroblasts as candidate effectors of FAO dysfunction in FCD and motivated direct functional interrogation.

To functionally validate our transcriptomic observations, we isolated and cultured primary intestinal fibroblasts from the terminal ileum of NIBD, non-FCD, and FCD patients. Under standard culture conditions, these cells spontaneously differentiated into myofibroblasts.(40)

Given our hypothesis that impaired FAO would lead to increased intracellular lipid accumulation, we assessed neutral lipid content using confocal imaging with the dye BODIPY 493/503. FCD-derived myofibroblasts exhibited markedly increased lipid droplet accumulation compared to non-FCD and NIBD controls. Flow cytometry of cells stained with BODIPY confirmed this increase in neutral lipid content, suggesting altered lipid handling.

To directly interrogate fatty acid utilization, we next examined mitochondrial fatty acid trafficking using a fluorescent pulse-chase assay(41). Cultured myofibroblasts were first incubated with BODIPY-C16, a fluorescent fatty acid analog, followed by mitochondrial labeling with PKmito Deep Red. Colocalization analysis revealed reduced overlap between fatty acid and mitochondrial signals in FCD-derived cells compared to non-FCD and NIBD controls. This decrease in colocalization suggests impaired trafficking of long-chain fatty acids into mitochondria for oxidation. Importantly, these defects were observed under standardized culture conditions, implying a persistent metabolic phenotype independent of exogenous inflammatory cues.

Consistent with these metabolic defects, we next evaluated whether impaired FAO in Crohn’s disease fibroblasts alters their responsiveness to PPARγ agonism during TGFβ-induced activation. Across all CD-derived fibroblasts (isolated from fibrotic or non-fibrotic lesions) examined, rosiglitazone showed markedly reduced ability to suppress TGFβ-induced αSMA expression compared to NIBD controls, indicating diminished PPARγ sensitivity. While our results focused on the overall CD versus NIBD distinction, internal heterogeneity was observed across CD samples with fibroblasts derived from fibrotic strictures displaying the least responsiveness to PPARγ agonism, suggesting that disease stage or prior in vivo conditioning may imprint a more refractory phenotype. These observations align with the concept of a stably reprogrammed, FAO-deficient myofibroblast state in CD and further support a model in which metabolic dysfunction contributes functionally to impaired resolution of fibroblast activation.

Reduced FAO has emerged as a unifying metabolic signature across fibrotic diseases involving the lung(10), liver(42), kidney(43), heart(44), and skin(45). In these contexts, the profibrotic cytokine transforming growth factor beta (TGFβ) plays a central role by driving fibroblast-to-myofibroblast differentiation and promoting myofibroblast activity, including extracellular matrix (ECM) deposition, contractility, and tissue remodeling.(46, 47) Mechanistically, TGFβ suppresses FAO in part through downregulation of PPARγ, a nuclear transcription factor essential for lipid metabolism and mitochondrial function.(35, 48) Loss of PPARγ activity impairs mitochondrial oxidative capacity, promotes intracellular lipid accumulation, and promotes differentiation to myofibroblasts and activation of the myofibroblast phenotype.(49, 50) In both in vitro and in vivo models of pulmonary(49), cardiac(51, 52), and dermal fibrosis(50, 53), PPARγ agonists have been shown to inhibit myofibroblast differentiation and ameliorate fibrotic remodeling.(29, 54) While this TGFβ–PPARγ–FAO regulatory axis is well-established in other organ systems, its relevance to intestinal fibrosis and CD, has remained largely unexplored. Our findings provide new evidence that this metabolic-fibrotic circuitry may also contribute to fibrostenotic progression in the context of CD.

Our data suggest that impaired FAO is not merely a consequence of fibrosis but may precede and contribute to its development. Early suppression of FAO-related genes in treatment-naive pediatric FCD samples points to metabolic reprogramming as a potential driver of fibrotic remodeling. The concomitant upregulation of glycolytic regulators suggests a metabolic shift away from oxidative metabolism, while loss of PPARγ activity and enhanced TGF-β signaling provide parallel routes to promote fibroblast activation. These findings suggest that once fibroblasts acquire a fully differentiated myofibroblast phenotype, they may become refractory to PPARγ-mediated metabolic rescue.

This observation aligns with the budding concept of “fibrotic memory,” wherein activated myofibroblasts maintain a stable transcriptional and mechanical phenotype even after the removal of profibrotic cues. Fibroblasts have been shown to retain “mechanical memory” that persists despite changes to exogenous cues, contributing to sustained activation.(40) Increasing evidence suggests that this memory is not solely mechanical but epigenetically reinforced. For example, studies in cardiac(55), kidney(56), and dermal fibroblasts(57) have demonstrated that persistent DNA methylation and altered chromatin accessibility maintain the expression of fibrotic genes even after the withdrawal of fibrotic stimuli such as TGFβ. Additionally, mechanical cues have also been shown to induce durable memory via histone modifications and noncoding RNAs, which may be reversible through targeted epigenetic therapies, such as histone deacetylase inhibitors (HDACi)(58). Consistent with this, preclinical studies have demonstrated encouraging antifibrotic potential of HDACi, although use has not extended to human trials yet.(59, 60) The persistent suppression of FAO in ex vivo FCD-derived myofibroblasts supports the existence of a stably reprogrammed fibrotic state in CD—potentially maintained through similar epigenetic mechanisms.

Together, these findings support the positioning of FAO impairment as a central metabolic signature of fibrotic myofibroblasts in CD. Clinically, this raises the possibility that FAO gene expression profiling in tissue or blood could serve as a biomarker for fibrotic risk. Notably, emerging epidemiologic evidence suggests that therapies influencing lipid metabolism may already modify disease course in IBD, as statin use has been associated with reduced progression to more severe disease phenotypes in a large nationwide cohort study(61). Moreover, early intervention to preserve FAO may prevent fibroblast activation and fibrosis progression. Although PPARγ agonists have demonstrated efficacy in preclinical models(49, 62–64), their translational success in human disease has been limited, highlighting the importance of therapeutic timing and disease stage.(60) Our findings also resonate with recent reports linking creeping fat and fibrostenotic progression in CD, which suggest that mesenteric fat-derived lipid flux may further interact with stromal metabolic reprogramming to promote stricturing disease(65). Strategies aimed at reversing fibrotic memory or restoring FAO in established myofibroblasts may offer new avenues for antifibrotic therapy in CD.

Our study has several limitations. While the use of primary human samples and integrative multi-omic analysis strengthens the validity of our conclusions, *in vitro* studies do not fully recapitulate the complex *in vivo* microenvironment. Fibroblast behavior is influenced by interactions with immune, epithelial, and endothelial cells, which were not captured in our study. Although our data suggests a role for FAO dysfunction in myofibroblast activation, the causal contribution of altered FAO to intestinal fibrosis requires validation in preclinical animal models. Future studies incorporating co-culture systems and *in vivo* perturbation of FAO pathways will be essential to determine the functional consequences of metabolic reprogramming in tissue remodeling and fibrosis progression.

In summary, our study identifies FAO dysfunction as a defining feature of fibrotic intestinal myofibroblasts in CD. Through a multi-omic approach integrating metabolomics, transcriptomics, and functional imaging, we uncover potentially targetable metabolic vulnerability in the fibrotic stroma. These findings lay the foundation for future translational studies aimed at intercepting intestinal fibrosis in CD via metabolic reprogramming.

## Methods

### Sex as a biological variable

This study included samples from both male and female human participants across all cohorts. Sex was recorded as a clinical variable and incorporated as a covariate in transcriptomic analyses to account for potential sex-related variability. The study was not designed to evaluate sex-specific biological effects, and results are therefore reported independent of sex. The findings are expected to be relevant to both males and females, as fibrostenotic Crohn’s disease affects both sexes and no prior evidence suggests sex-restricted metabolic mechanisms in intestinal fibrosis.

### Patient Cohorts and Clinical Phenotyping

Adult and pediatric patients with Crohn’s disease (CD) and non-inflammatory bowel disease (NIBD) controls were recruited from the University of North Carolina Hospitals and included in this study, which was approved by the UNC Institutional Review Board (19–2519, 15–0024, 10–0355, 17–0236). Clinical phenotyping captured demographic and clinical variables including age, sex, disease duration, age at diagnosis, age at sample acquisition, disease location, and disease behavior. Summary characteristics of the metabolomic cohort are shown in Table 1, and detailed patient-level clinical characteristics are provided in Supplementary Table 1. The adult and pediatric RNA-sequencing cohorts have been previously described (66).

### Metabolomics and Lipidomics

All sample preparation, data acquisition, and analysis for metabolomics and lipidomics was performed by Metabolon, Inc (Research Triangle Park, NC) per manufacture’s established sample preparation, UPLC-MS/M Data acquisition and analysis protocols.

### Multivariate and pathway level analyses of metabolomic and lipidomic data

Sparse partial least squares discriminant analysis (sPLS-DA) regression was performed on imputed, log-transformed, median-scaled metabolite values from both the HD4 and Complex Lipid Panel (CLP) assays, using the splsda() function from the mixOmics (v6.22.0) R package (67). The optimal number of components (“ncomp”) and variables (“keepX”) to include in each sPLSA-DA model were determined using the tune.splsda() function (2 components for the HD4 dataset, and 3 components for the CLP dataset). Results of the plotIndiv() and plotVar() functions, for sample and correlation plots respectively, were plotted using ggplot2 (v3.4.4) (68). Lollipop plots of the acylcarnitines and complex lipids of interest were generated using metabolomics results of the HD4 and Complex Lipid Panel (CLP) assays. Fold change values (also referenced in the figure legends as “Enrichment”) are the ratio of mean scaled metabolite values of CD samples relative to NIBD samples. Statistics were reported by Metabolon for each comparison derived from the natural log-transformed data. A Student’s t-test was performed for each comparison, and the resulting p-values were log-transformed for plotting. Plots were created with the ggplot2 (v3.4.4) R package (68).

### Cell culture

All cells were cultured at 37°C in a humidified incubator with 5% CO₂. Medium was changed every 2–3 days, and fibroblasts were passaged at a 1:3 ratio upon reaching 80% confluence.

Cells were regularly tested and confirmed to be mycoplasma-free using Mycostrip tests (Invivogen).

### Primary human fibroblast isolation and culture

Characteristics of donors from whom primary cells are isolated are described in Table 2. Primary human intestinal fibroblasts were isolated from mucosal tissue using enzymatic digestion in 30 mL of pre-warmed EDTA-HBSS at 37°C for 30 minutes with shaking at 250 rpm. Following this, dithiothreitol (DTT) was added (final concentration 1 mM), and the tissue was incubated for an additional 10 minutes. Mucosal fragments were then collected and transferred into 20 mL of pre-warmed collagenase-HBSS, composed of HBSS (with Ca²⁺/Mg²⁺), 2.5% heat-inactivated fetal bovine serum (FBS), 1% penicillin/streptomycin (P/S), 1 mg/mL Collagenase IV (Gibco, Cat# 17104019), and 0.1 mg/mL DNase I (Roche, Cat# 10-104-159-001). Samples were incubated at 37°C for 30 minutes with shaking at 250 rpm. The resulting cell suspension was filtered through a 70 µm cell strainer to remove debris. Cells were pelleted by centrifugation at 2500 rpm for 5 minutes, rinsed with phosphate-buffered saline (PBS), and resuspended in 10 mL of complete DMEM/F12 medium supplemented with 10% FBS and 1% P/S. To assess chromosomal stability of primary cells, karyotype analysis was performed by KaryoLogic (Durham, NC). Aneuploidy was detected by passage 13; therefore, cells beyond passage 13 were retired from culture and excluded from experimental analyses. Culture purity and differentiation to myofibroblasts were confirmed through flow cytometry (see section below).

**TABLE 2:** Donor sample characteristics.

| ID | Disease | Collection procedure |
| --- | --- | --- |
| CHAMP80 | NIBD | Surgical |
| CHAMP81 | NIBD | Surgical |
| CHAMP 83 | NIBD | Surgical |
| CHAMP142 | NIBD | Surgical |
| CHAMP61 | CD - Fibrotic | Surgical |
| CHAMP66 | CD- Fibrotic | Surgical |
| CHAMP74 | CD- Fibrotic | Surgical |
| CHAMP97 | CD- Fibrotic | Surgical |
| CHAMP70 | CD – Non-fibrotic | Surgical |
| 17-0236-0619 | CD – Non-fibrotic | Colonoscopy |
| 17-0236-0625 | CD – Non-fibrotic | Colonoscopy |
| 17-0236-0637 | CD – Non-fibrotic | Colonoscopy |

### Cell lines

CCD-18Co (ATCC CRL-1459), a human colonic fibroblast cell line was cultured in Eagle’s Minimum Essential Medium (EMEM) supplemented with 10% heat-inactivated FBS and 1% P/S, according to ATCC recommendations.

Lymphoblastoid cell line (LCLs; gift of Dr. Blossom Damania, UNC Chapel Hill) was generated by purifying human peripheral blood for CD19+ B cells subsequently infected with Epstein-Barr virus (MOI = 0.1). LCLs were maintained in RPMI 1640 medium (Gibco) supplemented with 10% FBS, 2 mM L-glutamine, and 1% P/S. LCLs are fast growing and accordingly were passaged at a 1:4 ratio.

### Cell-based assays

Cell density for all cell-based assays was optimized at 1E5 cells/ml. Cells were allowed to adhere to culture vessels overnight prior to exposure to (a) rosiglitazone or vehicle controls for 24-72 hours to quantify cell viability using CellTiterGLO assay (Promega) following the manufacturer’s instructions; or (b) rosiglitazone and/or TGFβ (10 ng/ml) for 48 hours, after which αSMA expression was assessed using western blots performed as previously described, by washing cells with ice-cold PBS and harvesting in lysis buffer containing 150 mM NaCl, 50 mM Tris⋅HCl (pH 8), 0.1% Nonidet P-40, 50 mM NaF, 30 mM β-glycerophosphate, 1 mM Na_3_VO_4_ and 1× Complete Protease Inhibitor Mixture (Roche). Proteins were electrophoresed on a 4-20% gradient polyacrylamide gel (BioRad) and transferred onto a Hybond-ECL nitrocellulose membrane (GE Healthcare). Membranes were blocked with 5% fat-free milk for 1 h and then incubated overnight at 4 °C with the indicated antibodies: αSMA (#14968), α/β Tubulin (#2148), and anti-rabbit IgG-HRP (#7074), all from Cell Signaling Technology. Bands were visualized by chemiluminescence on iBright imager (Invitrogen). Images were minimally processed with Photoshop to adjust sizing.

### Flow cytometry and analysis

Culture purity and differentiation into myofibroblasts were assessed by flow cytometry. Cells were stained with Live/Dead Fixable Blue Dead Cell Stain (Invitrogen, Cat #L23105) in PBS at 4°C for 30 minutes to exclude non-viable cells. Following a PBS wash, surface staining was performed using CD90 BV605 (BD, Cat #747750) to identify fibroblasts. A dump channel strategy was employed using a cocktail of CD326 APC (Invitrogen, Cat #MA538715), CD31 APC (Invitrogen, Cat #17-0319-42), and CD45 APC (Proteintech, Cat #APC-65109100TST) to exclude epithelial, endothelial, and hematopoietic cell contaminants, respectively.

Cells were then fixed and permeabilized using the BD Cytofix/Cytoperm Kit (BD Biosciences) for 20 minutes at room temperature, followed by intracellular staining with α-smooth muscle actin (αSMA) AF488 (Invitrogen, Cat #53-9760-82).

For all flow cytometry experiments, single-stained compensation controls were prepared using UltraComp eBeads Plus Compensation Beads (Invitrogen). A lymphoblastoid cell line (LCL), negative for both CD90 and αSMA, was included in all experiments as a biological negative control. Data were acquired on a BD LSRFortessa flow cytometer and analyzed using FlowJo software (BD Biosciences). Statistical analyses were performed using GraphPad Prism.

For neutral lipid staining, cells were incubated with 5 μM BODIPY 493/503 (Life Technologies) in PBS at 37°C for 30 minutes prior to fixation. Surface staining for CD90 was performed as above. Cells were then fixed and permeabilized as previously described, followed by intracellular staining for αSMA APC (R&D Systems, Cat #IC1420A). After gating on CD90⁺ αSMA⁺ cells, mean fluorescence intensity (MFI) of the BODIPY signal was quantified using FlowJo.

### Bulk RNA-sequencing and analysis

#### Adult colonic cohort

Total RNA was extracted from primarily colonic mucosa samples for paired-end RNA-sequencing. Reads were aligned to the human genome (hg38/GRCh38) based on GENCODE v47 annotations using STAR (v2.7.9a). To prevent allelic mapping biases, reads were aligned using the ‘’--WaspOutputModè’ with corresponding sample genotypes when running STAR. We obtained 108 uninflamed colonic CD tissue and 58 uninflamed non IBD control samples. Genes were filtered to retain those with at least five counts in more than twenty five percent of samples.(69) Gene annotation and identifier mapping were performed using the Bioconductor org.Hs.eg.db package.(70) Sex, batch detail, and transcript integrity number were encoded as covariates, with transcript integrity number scaled to mean zero and unit variance. To estimate empirical negative control genes for removal of unwanted variation, we first fit a simple DESeq2 model with design equal to disease, using size factor estimation with positive counts and without independent filtering.(71) From this model we extracted log2 fold changes and p values for Crohn disease versus non inflammatory bowel disease. The variance stabilizing transformation was then applied to the fitted object to obtain a variance stabilized expression matrix, and per gene variance was computed with matrixStats. Candidate negative control genes were defined using four criteria. First, they showed little evidence of differential expression for disease, with absolute log2 fold change less than or equal to 0.25 and p value at least 0.5. Second, they were in the lowest twenty percent of the variance distribution across all samples. Third, they had weak correlation with nuisance covariates. To quantify this, we formed a covariate matrix containing log transformed library size, scaled transcript integrity number, sex, batch detail, and a numeric coding of disease, and computed for each gene the maximum absolute Pearson correlation between variance stabilized expression and any covariate. Genes with maximum absolute correlation less than or equal to 0.2 were retained. Fourth, genes mapped to chromosomes X and Y were excluded from the negative control pool using an external annotation table. The intersection of these filters yielded a pool of empirical 324 control genes that were selected as negative controls for RUVSeq.

For differential expression in the adult colon, we restricted the dataset to non-inflammatory bowel disease (NIBD) and Crohn disease samples and fit a DESeq2 model with design equal to batch detail, sex, scaled transcript integrity number, all 5 RUV factors, and disease.(72) Disease was modeled with NIBD bowel disease as the reference. Wald tests were used to estimate log2 fold changes and Benjamini Hochberg adjusted p values for Crohn disease versus non inflammatory bowel disease.

To summarize pathway level transcriptional patterns, we defined an a priori gene panel that grouped genes into fatty acid oxidation, PPAR gamma axis, glycolysis, and transforming growth factor and extracellular matrix categories. For each gene in this panel, we extracted the DESeq2 log2 fold change for Crohn disease versus non inflammatory bowel disease and assembled a two-column matrix in which non inflammatory bowel disease was set to zero and the Crohn disease column contained the estimated log2 fold change. This matrix was visualized using ComplexHeatmap with a fixed color scale and row annotation specifying pathway membership.(73)

*Adult ileal resection cohort:* Total RNA was extracted from primarily ileal mucosa samples for paired-end RNA-sequencing. Reads were aligned to the human genome (hg38/GRCh38) based on GENCODE v47 annotations using STAR (v2.7.9a). To prevent allelic mapping biases, reads were aligned using the ‘‘--WaspOutputModè‘ with corresponding sample genotypes when running STAR. We obtained 74 inflamed ileal CD samples and 26 ileal non IBD samples. From the CD ileal samples, 24 samples were labeled as no ileal stricturing while 50 samples we labeled as ileal stricturing present at the time of sample collection. Adult ileal RNA sequencing data were processed in an identical framework to adult colonic cohort. 218 genes were selected as negative control for RUVseq. Raw counts and metadata were passed to RUVg with k equal to five to obtain latent factors W1 through W5. To compare non inflammatory and fibrostenotic phenotypes within Crohn disease, we defined a three level Group variable with levels non inflammatory bowel disease, Crohn disease non fibrostenotic, and Crohn disease fibrostenotic based on ileal stricture status at time of sample collection. A DESeq2 model was fit with design equal to sex, transcript integrity number, all 5 RUV factors, and Group, using non inflammatory bowel disease as the reference. Contrasts were extracted for Crohn disease non fibrostenotic versus non inflammatory bowel disease and Crohn disease fibrostenotic versus non inflammatory bowel disease. Using the same fatty acid oxidation, PPAR gamma axis, glycolysis, and transforming growth factor and extracellular matrix gene panel defined above, we assembled a three-column matrix of log2 fold changes with non-inflammatory bowel disease set to zero and the two Crohn disease groups represented by their respective log2 fold changes.

*Treatment naive pediatric ileal cohort:* For the pediatric ileal cohort, we obtained 45 non-IBD controls with a total of 43 CD patients as previously described(66). 11 of the CD samples were labeled as FCD as they developed stricturing in longitudinal follow-ups while 30 were labeled as non-FCD as they did not develop strictures.

Average expression for each gene across all samples using the variance stabilized counts and selected the five thousand most highly expressed genes was performed. From this subset we calculated the standard deviation divided by the mean expression for each gene and chose the one thousand genes with the lowest relative variability as candidate negative controls. Counts for these genes were rescaled and rounded, and a new SeqExpressionSet was created. RUVg was then run with k equal to one, using these genes as negative controls, to estimate a single latent factor W1. A new DESeq2 object was then constructed using the full count matrix and the augmented metadata with design equal to batch, sex, transcript integrity number, W1, and a three level fibrostenotic status variable encoding non inflammatory bowel disease, non fibrostenotic Crohn disease, and fibrostenotic Crohn disease. Lowly expressed genes were removed with the edgeR filterByExpr function before model fitting.(74) For panel level visualization, we focused on contrasts that used non inflammatory bowel disease as the reference. From the contrast results we extracted log2 fold changes for the prespecified gene panel spanning fatty acid oxidation, PPAR gamma axis, glycolysis, and transforming growth factor and extracellular matrix genes.

#### Single-cell RNA-sequencing analysis of stromal gene expression in Crohn’s disease ileum and healthy colon

Previously processed stromal single-cell RNA-sequencing data from(15) were loaded into R (v4.4.2) and analyzed using the Seurat package (v5.3.0)(75). Cells were subset to fibroblasts based on the original cell type annotations provided in the published dataset. For visualization, a quantitative dot plot was generated using all fibroblast-annotated cells, displaying representative genes associated with glycolysis, myofibroblast activation, inflammation, PPARγ signaling, TGF-β signaling, and fatty acid oxidation (FAO). Gene sets for pathway classification were derived from a combination of manual annotation and curated databases. Pathway gene lists were downloaded from the Molecular Signatures Database (MSigDB)(76) and included BioCarta pathways for PPARγ and TGF-β signaling, as well as Gene Ontology (GO)(77) annotations for fatty acid oxidation (GO:0019395 – “Fatty Acid Oxidation”) and glycolysis (GO:0006110 – “Regulation of Glycolytic Process”). Pseudobulk differential expression analysis was performed by aggregating raw counts from fibroblast-annotated cells within each tissue compartment (normal colon, non-inflamed ileum, inflamed ileum, and strictured ileum) for each patient.

Analyses were restricted to pseudobulk groups containing at least 50 fibroblast cells per sample. Differential testing was performed using DESeq2 (v1.46.0)(72), installed through Bioconductor (v3.20)(78).Genes with an adjusted *P* ≤ 0.1 (Benjamini–Hochberg false discovery rate) were considered significant. Heatmaps displaying log₂ fold-change values for significantly altered genes were generated using TidyHeatmap (v1.12.2)(79).

#### Confocal microscopy

For lipid droplet quantification, human intestinal myofibroblasts were seeded at 1,000 cells per well into 8-well chambered coverglass slides (Cellvis, Cat# C8-1.5P) and allowed to adhere overnight under standard culture conditions. The following day, cells were fixed with 4% paraformaldehyde for 10 minutes at room temperature and washed twice with phosphate-buffered saline (PBS). Fixed cells were incubated with 1 μM BODIPY 493/503 (Life Technologies) in PBS for 30 minutes to stain neutral lipids. Nuclei were counterstained with Hoechst 33342. Lipid droplet quantification was performed by manual droplet counting and normalized to cytoplasmic area using Fiji (ImageJ).

For mitochondrial fatty acid trafficking and colocalization analysis, human intestinal myofibroblasts were seeded as described above. After overnight adherence, cells were incubated with 25 nM C16 BODIPY (Invitrogen, Cat# D3821) in complete growth media for 16 hours to label intracellular fatty acid pools. Following incubation, cells were washed with PBS and subjected to a 6-hour chase period in fresh complete media to allow for fatty acid mobilization. Mitochondria were subsequently stained with PKmito Deep Red (Spirochrome, Cat# CY-SC055) at a 1:10,000 dilution in complete media, and nuclei were counterstained with Hoechst 33342.

Prior to imaging, cells were washed with PBS to remove excess dyes. Both fixed and live-cell confocal imaging were performed using a Zeiss LSM 880 confocal microscope equipped with ZEN acquisition software (Zeiss). Colocalization analysis was performed using Fiji (ImageJ) with the BIOP JACoP plugin to calculate both Pearson’s correlation coefficient and Manders’ overlap coefficient. Image thresholding for colocalization analysis was performed using Otsu’s method for both the C16 BODIPY and PKmito channels to minimize background and ensure consistent segmentation.

### Statistical analysis

Statistical testing was performed using GraphPad Prism 10.2.0 software (GraphPad, San Diego, CA). Sample size and specific analysis explained in figure legends. P-values below 0.05 are considered significant and indicated within the figures and in-text.

## Author Contributions

J.A., A.P.B., and S.Z.S. conceived the study. J.A. performed experiments, analyzed and interpreted data, and wrote the manuscript.

A.A., B.D.M., V.G., and V.T. performed experiments, analyzed data, generated figures, and contributed to manuscript writing and editing.

B.H. and D.W. provided technical support, assisted with sample acquisition, and contributed to figure and table generation.

G.L., C.B., G.W.-J.L., and S.S. assisted with sample acquisition and provided technical support.

M.K. contributed to patient sample acquisition.

A.C.S. contributed to critical revision and editing of the manuscript.

F.R. provided data and contributed to manuscript editing.

J.E.T. provided guidance on experimental design and contributed to manuscript editing.

T.S.F. supervised data analysis and provided intellectual input.

A.P.B. and S.Z.S. provided funding, supervised the study, and critically revised the manuscript. All authors reviewed and approved the final manuscript.

## Patient and public involvement

Patients and the public were not involved in the design, conduct, reporting or dissemination plans of this research.

## Ethics approval

This study involved human participants and was approved by the Institutional Review Boards at the University of North Carolina at Chapel Hill (IRB numbers: 19-2519, 15-0024, 10-0355, 17-0236). Written informed consent was obtained from participants or was waived where appropriate for the use of de-identified surgical specimens, in accordance with institutional guidelines. This study did not involve animal subjects.

## Acknowledgements

The authors acknowledge Wendy Salmon, Director of the UNC Hooker Imaging Core, for expert guidance in optimizing confocal co-localization imaging experiments. Flow cytometry studies were performed with support from the UNC Flow Cytometry Core Facility.

---

**Supplementary Figure 1.**
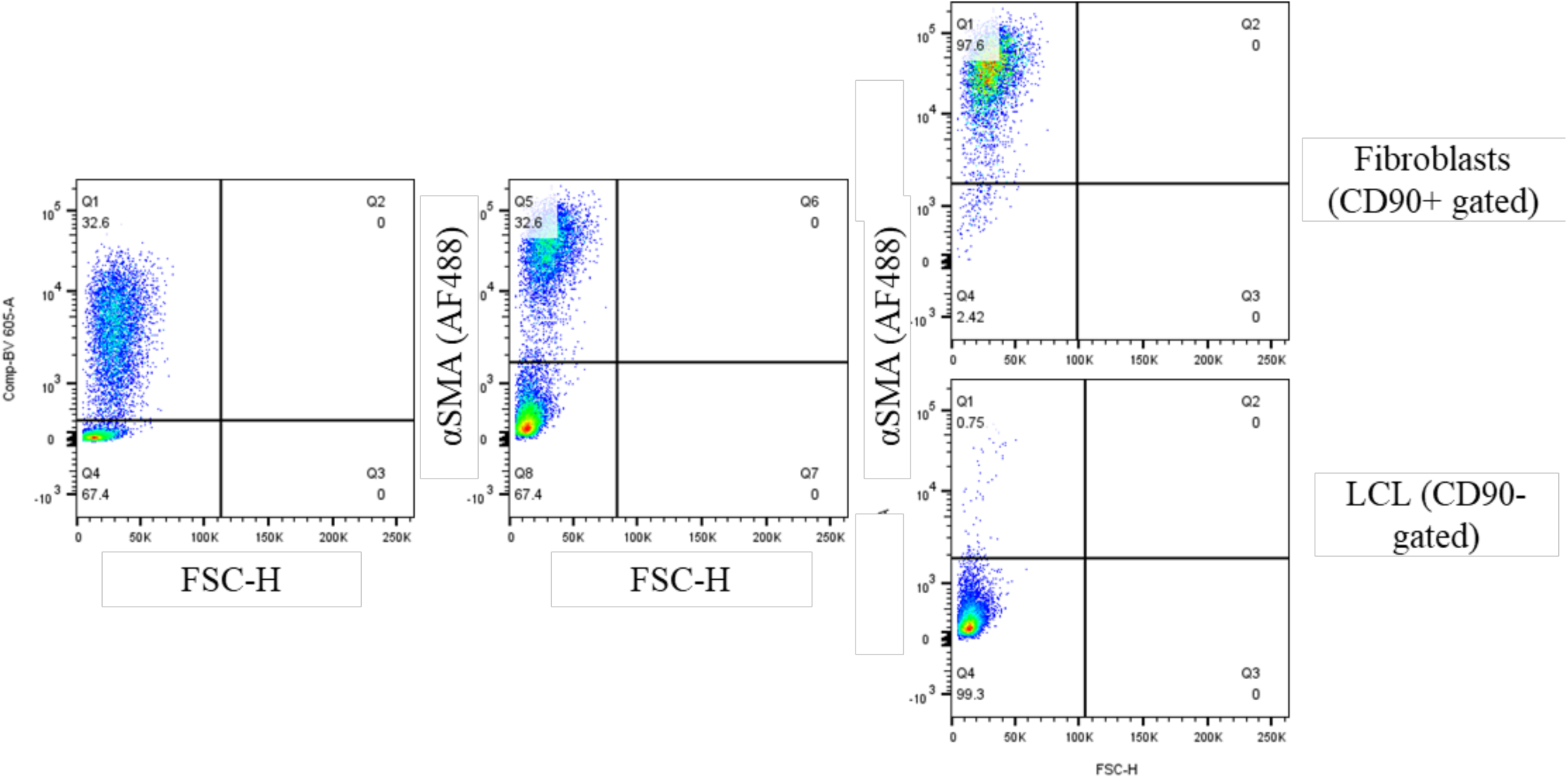
Gating strategy for isolation and validation of primary intestinal myofibroblasts by flow cytometry. Primary intestinal myofibroblasts were identified by CD90 (Thy-1) expression and validated using αSMA staining. Singlet and viability gating (not shown) were applied before CD90-based selection. A lymphoblastoid cell line (LCL) was included as a CD90⁻/αSMA⁻ negative controls. Left panels show mixed-cell populations with CD90 and αSMA staining. CD90⁺-gated cells showed uniform αSMA positivity, confirming enrichment for myofibroblasts, while CD90⁻-gated LCL control cells remained αSMA⁻. Representative plots from a non-FCD (B1) Crohn’s disease sample demonstrate clear separation between fibroblast and non-fibroblast populations and validate myofibroblast identity for downstream assays.

**Supplementary Figure 2.**
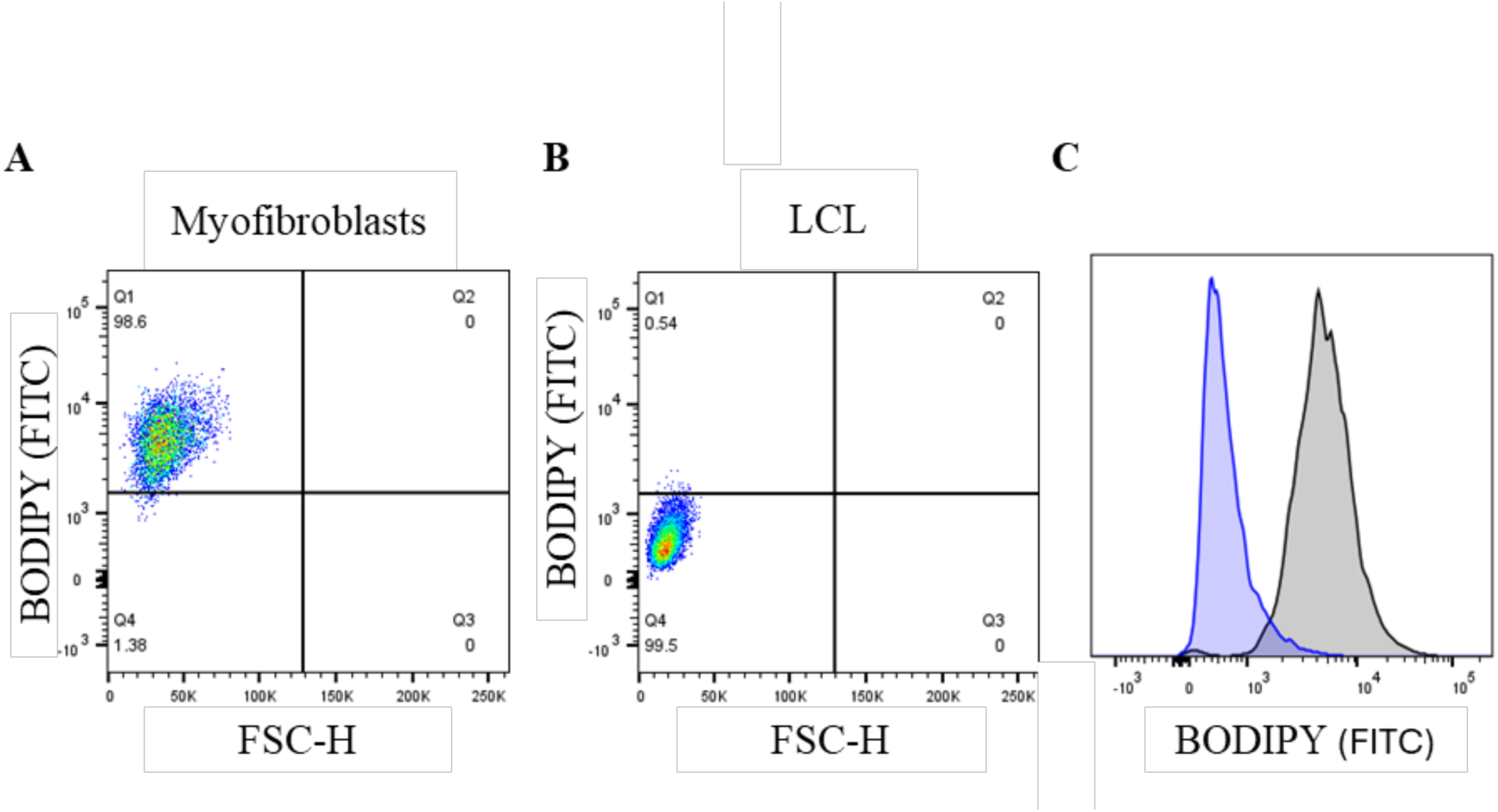
Flow cytometry strategy for quantifying neutral lipid content in primary intestinal myofibroblasts using BODIPY 493/503 staining. **(A)** Primary intestinal myofibroblasts were stained with BODIPY 493/503 to assess neutral lipid content. Standard singlet and viability gating (not shown) were applied before measurement. **(B)** A lymphoblastoid cell line (LCL) was included as an unstained negative control to establish background FITC fluorescence. All samples were acquired using identical PMT settings. Representative dot plots from a non-FCD (B1) Crohn’s disease sample demonstrate high BODIPY signal in myofibroblasts compared with minimal autofluorescence in LCL controls. **(C)** Corresponding histograms show clear separation of FITC intensity between BODIPY⁺ myofibroblasts (gray) and the LCL negative control (blue), validating the specificity of the lipid-staining assay.

**Supplementary Figure 3.**
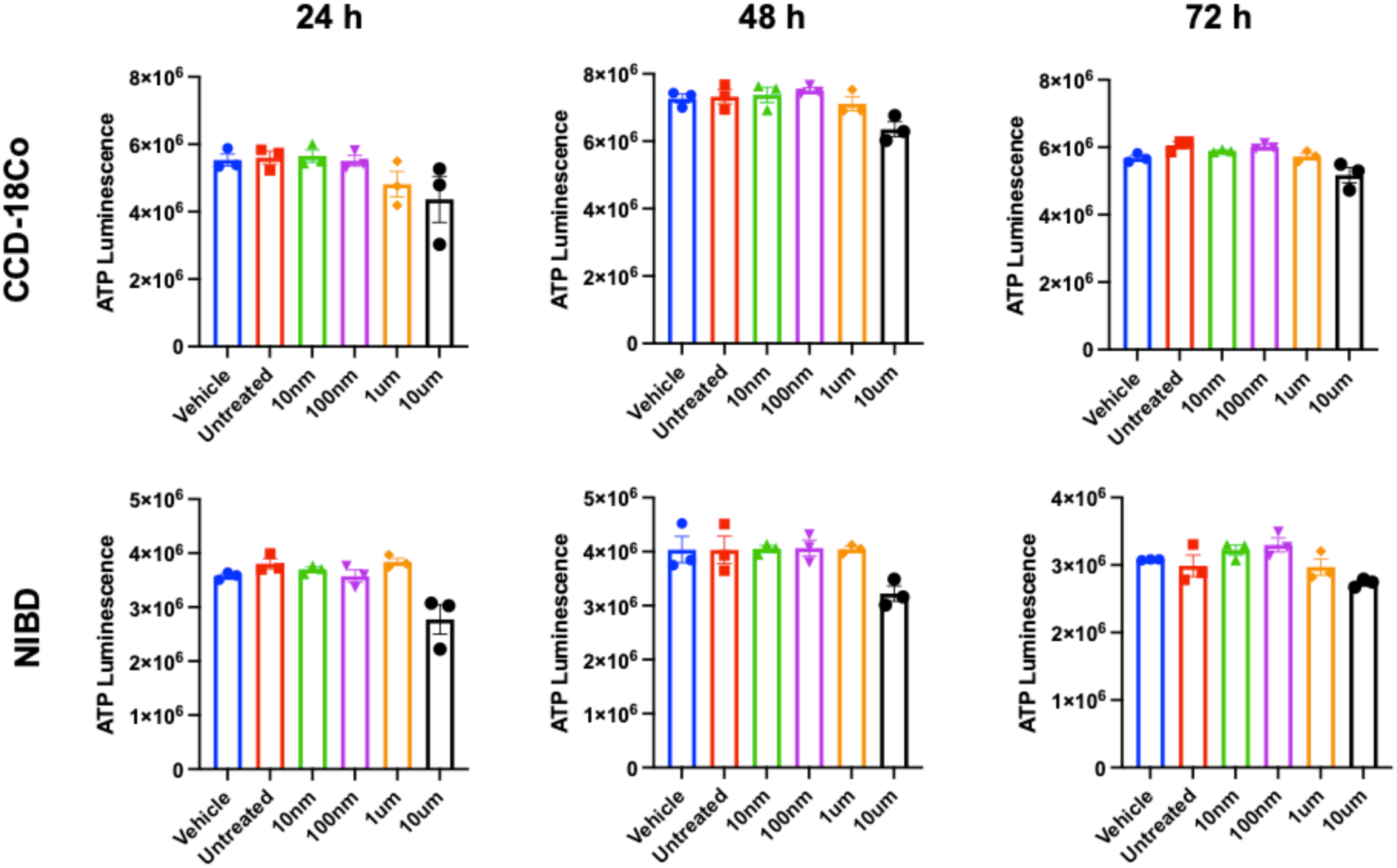
Rosiglitazone does not impair cell viability in CCD-18Co or primary fibroblasts. CCD18Co (top row) or primary (NIBD; bottom row) fibroblasts were plated in 96 well plates at 1E5 cells/ml and treated with increasing concentrations of rosiglitazone (10 nM–100 µM) or vehicle (0.1% DMSO) for 24, 48, or 72 hours. Cell viability was assessed by CellTiter GLO ATP luminescence assay. No statistically significant reduction in normalized ATP luminescence was observed at any dose or timepoint, confirming lack of drug toxicity under the experimental conditions. Data represent mean ± SEM of three replicates.

